# Epigenetic alterations in gut and brain of adult rats after oral administration of miR-320-3p and miR-375-3p at mid-lactation, and preventive potential of miR-320-3p on early weaning stress

**DOI:** 10.1101/2022.06.20.496755

**Authors:** Gabriel A Tavares, Amada Torres, Gwenola Le Drean, Maïwenn Queignec, Blandine Castellano, Laurent Tesson, Séverine Remy, Ignacio Annegone, Sandra L de Souza, Bruno Pitard, Bertrand Kaeffer

## Abstract

**Aim:** To investigate if the artificial delivery of microRNAs naturally present in the breastmilk can impact the gut and brain of young rats according to weaning.

**Methods:** Animals from a new transgenic rat line expressing green-fluorescent protein in the endocrine lineage (cholecystokinin expressing cells) received at Day-12, near neural diversification, a single oral bolus of mir-320-3p or miR-375-3p, embedded in DiOleyl-Succinyl-Paromomycin (DOSP), and were further early (Day-15) or regularly (Day-30) weaned. Relevant miRNA (miR-320-3p, miR-375-3p, miR-375-5p, miR-16-5p, miR-132-3p, miR-504), *polr3d*, *hspb6*, inflammation, enteroendocrine, and circadian clock-related mRNAs, chromatin complexes, and duodenal cell density were assayed at 8h post-inoculation and at Day-45.

**Results:** The miR-320-3p/DOSP induced immediate effects on H3K4me3 chromatin complexes with polr3d promoter (p<0.05) but no long-term effects. On regular weaning, at Day-45, both miR-320-3p and 375-3p were down-regulated in the stomach, up-regulated in the hypothalamus (p<0.001) but only miR-320-3p was up-regulated in the duodenum. After early weaning, the miR-320-3p and miR-375-3p levels were down-regulated in the stomach and the duodenum, but up-regulated in the hypothalamus and the hippocampus. Combining miR-320-3p/DOSP with early weaning enhanced miR-320-3p and chromogranin A expression in the duodenum. In the hippocampus, the miR-504 was down-regulated for both sexes, but in the brain stem, up regulated only for females, along with miR-320-3p and miR-16-5p levels. In the hypothalamus, clock levels were up regulated for both sexes. In the miR-375-3p/DOSP group, the density of enteroendocrine duodenal cells increased. The long-term effect of miR-375-3p/DOSP was more limited, according to the fourfold lower number of predicted targets than with miR-320-3p.

**Conclusion:** Addressing oral miRNA-320-3p loads to duodenal cell lineage is paving the way for the design of new therapeutics, manipulating long term consequences of early life stress.

## Introduction

In recent years considerable evidence has demonstrated that the adult health status may be strongly influenced by experiences in early life modulated by epigenetic changes.^1, 2^ The inadequate interruption of lactation (Early weaning) impairs an important nutritional and maternal contact, promoting anxiety, depression, or stress in neonates, some with deleterious lifelong consequences. The miR-504 and miR-16-5p have been identified as key miRNAs, in the relationship between early life stress and the modulation of the dopaminergic and serotonergic systems.^3^ The miR-504 directly targets the 3′UTR of the dopamine D1 receptor gene (*drd1*)^4^ and the miR-16-5p is involved in the regulation of the serotonin transporter (*sert*) in the raphe of depressive rats.^5^ Moreover, the miR-132-3p has been described in neural cell epigenetics,^6^ coupling circadian rhythms and the daily rhythms of neuron plasticity involved in cognition.^7^ Although only a few changes in miRNA expression were reported after maternal separation, these studies support the notion that Early Life Stress induces susceptibility to later life stress at the epigenome level.

Moreover, the absorption of miRNAs in the rat stomach has been demonstrated to be under the dependence of Systemic RNA interference–deficient transporter (SIDT1)^8^, opening the possibility of the natural transit of miRNAs present in breast milk from mother to offspring. Consequently, manipulating the physiopathology of rat pups through the use of extracellular miRNAs, given as supplements, will integrate new knowledge for preventing at the earliest time possible the onset of chronic pathology. The oral delivery of extracellular miRNA is of general interest^9, 10^, but we have little knowledge on the immediate and long-term effects on the molecular phenotype of a model organism, as well as on the consequences of combining with early-life stress. Milk contains a high amount of miRNAs which has been proposed for transfer between mother and child for immune regulation^11^, priming the immune system of the lactating infant when from plant origin^12^, transgenerational health influence^13^, or for trans-species effect on adult consumers through dairy products.^14, 15^ Here, we have focused on two microRNA common to rat and human breast milk and used for their delivery previously developed lipidic derivatives of natural aminoglycosides, allowed in food, shown to be efficient for intracellular delivery of siRNA, DNA, mRNA, or miRNA.^16–21^ The miR-320-3p is associated to breast milk exosomes^22^ and a highly conserved miR among mammals.^23^ A *cis*-regulatory role is known for miR-320-3p which participates in a negative feedback loop at the polr3d promoter inducing transcriptional gene silencing in Human Embryonic Kidney-293 cells.^24^ In rat pups, we have reported an immediate effect of miR-320-3p administered orally on hspb6, polr3d mRNAs and polr3d promoter in chromatin complexes.^20^ The miR-320-3p is known for its bioactivity in various diseases from type-2 diabetes, and inflammatory bowel disease, to atherosclerosis.^25–28^ The miR-375-3p is one of the most abundant miRNA in the gastrointestinal tract, impacting the homeostasis of the enteroendocrine lineage of mucosal cells.^29^ It has been related to child depression^30^, and the differentiation of mouse neurites in the hippocampus.^31^ This miRNA may be involved in neuroprotective mechanisms in response to stress^32, 33^ and has further been associated with Alzheimer’s Disease.^34^

In this paper, we demonstrate that force-feeding with miR-320-3p/DOSP at mid-lactation induced long-term effects on gastrointestinal epithelium and brain of young rats, deeply altering the regulation of endogenous miR-320-3p and miR-375-3p.

## Material and methods

### 1. Study design (Figure 1)

As shown in Figure 1A, we have followed the immediate effect of miRNA bolus on rat pups at Day-12 at the beginning of the dark phase and 8 hours after oral administration.^20^ Here, we apply the miRNA supplementation from Day-12 of age because extensive changes in gene expression of neurodevelopmental processes related to cell differentiation and cytoskeleton organization, have been identified in the hypothalamus of rat pups born from low protein-fed mothers.^35^ Rats were sacrificed at Day-45 to evaluate long-term effects on the physiology^36^ after force-feeding with either miR-320-3p or miR-375-3p embedded in DOSP. The control groups were force-fed with the vehicule solution of DOSP. The rats pre-treated with miR-320-3p/DOSP or corresponding control groups endured or not an early weaning, before sacrifice at Day-45. The rat pre-treated with miR-375-3p/DOSP received a regular weaning. The miR-320-3p/DOSP and miR-375-3p/DOSP groups, both enduring a regular weaning, were used for evaluation of the long-term effects of 2 unrelated miRNAs. We have used 2 weaning times at Day-15 (Early) or Day-30 (Regular) applied after the single oral bolus of miR-320-3p/DOSP in order to compare consequences of the treatment combined with early life stress against stress-less weaning.^37^ Rat pups separated from their mothers at D-15 were fed on soup made from standard chow. Litters were maintained at the UMR-1280 husbandry, allocated in rooms either with a light on at 7:00 am or off at 7:00 pm. Our experimental protocol was approved by the “Comité d’éthique pour l’expérimentation animale, Pays de la Loire, France” under number #APAFIS-21917. Studies on rats were performed according to the rules of the Nantes animal experimental unit [in compliance with the European Communities Directive of 2010/63/UE, 22 September 2010]. The total number of mothers was of 9. The first litter with 12 rat pups was used for immediate effect of miR-320-3p/DOSP, miR-375-3p/DOSP, or control made up of DMEM/PBS0 (4 pups per combination at random, sacrifice at 8 Hours). Eight litters were balanced at birth to 8 rat pups (4 males, 4 females). For oral inoculation, all solutions of deep-frozen miRNAs and of DOSP kept at 4°C, were warmed to room temperature. Rat pups were maintained at warm (37°C) in a parallel transparent box next to the mother box during one hour for gastric emptying. Mixtures of miRNAs and DOSP were extemporaneously prepared. Force-feeding was gently applied by trained experimenters allowing the delivery of 9 x 10^8^ molecules of miRNA in the stomach at the beginning of the dark phase.^20^ Gastric fluids were collected to evaluate a putative re-export of loaded miRNA though extracellular vesicles. The stomach cell wall was sampled as the inoculation site and the duodenum was relevant for our transgene expression. All q-PCR analyses were done on 8 rats, when we did not found a sex effect. The rationale for choosing brain area was for the hypothalamus as the main center of the major center of energy homeostasis, the hippocampus as involved in the memory of food reward and choice, the brain stem as the outcome of the vague nerve linking the intestine to the brain. We had a strong sex effect in brain, so we did all q-PCR analyse on 4 rats per sex. The effects have been screened according to gene sets related to inflammatory and enteroendocrine status of stomach or duodenum, and on the serotoninergic/dopaminergic balance of brain areas. Our transgenic GFP-CCK-p rat derived from the Sprague-Dawley strain is allowing to follow an enteroendocrine cell lineage labeled with Green Fluorescent Protein (GFP) in duodenum crypts.

**Figure 1.**
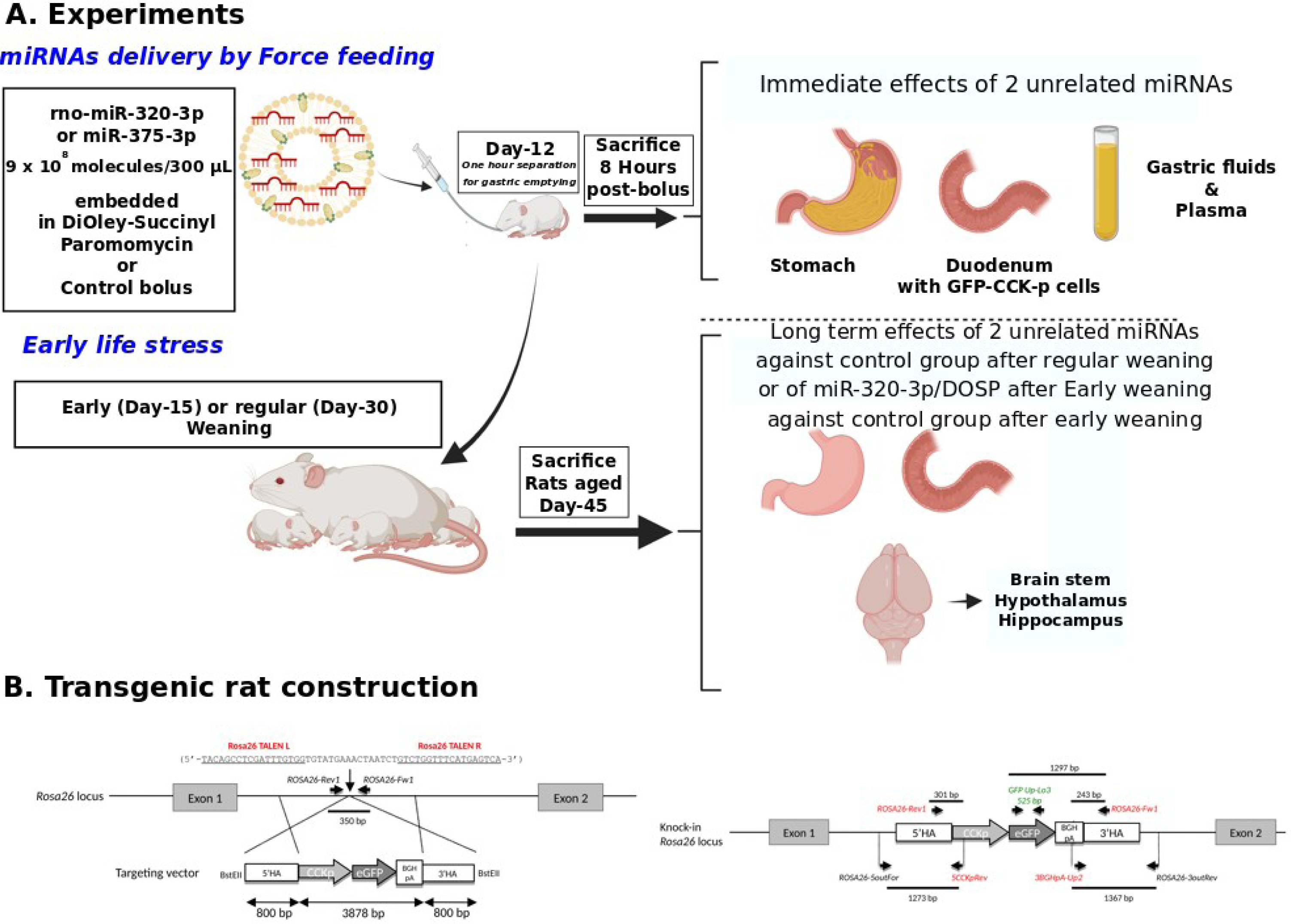
Study design. A. Experiments. we have followed the immediate effect of miRNA bolus on rat pups at Day-12 at the beginning of the dark phase and 8 hours after oral administration.^20^ We have sacrificed rats at Day-45 to evaluate long-term effects on the physiology^36^ of rats force-fed with either miR-320-3p or miR-375-3p embedded in DOSP against controls force-fed with the vehicule solution of DOSP. The rats pre-treated with miR-320-3p/DOSP endured or not an early weaning, before sacrifice at Day-45. The rat pre-treated with miR-375-3p/DOSP received a regular weaning, and sacrificed at Day-45. The miR-320-3p/DOSP and miR-375-3p/DOSP were used for evaluation of the long-term effects of 2 unrelated miRNAs. We have used 2 weaning times at Day-15 (Early) or Day-30 (Regular) applied after the single oral bolus of miR-320-3p/DOSP in order to compare midterm consequences of the treatment combined with early life stress against stress-less weaning.^37^ Rat pups separated from their mothers at D-15 were fed on soup made from standard chow. Litters were maintained at the UMR-1280 husbandry, allocated in a room either with a light on at 7:00 am or off at 7:00 pm. Our experimental protocol was approved by the “Comité d’éthique pour l’expérimentation animale, Pays de la Loire, France” under number #APAFIS-21917. Studies on rats were performed according to the rules of the Nantes animal experimental unit [in compliance with the European Communities Directive of 2010/63/UE, 22 September 2010]. The total number of mothers was of 9. The first litter with 12 rat pups was used for immediate effect of miR-320-3p/DOSP, miR-375-3p/DOSP, or control made up of DMEM/PBS0 (4 pups per combination at random, sacrifice at 8 Hours). Eight litters were balanced at birth to 8 rat pups (4 males, 4 females). For oral inoculation, all solutions of deep-frozen miRNAs and of DOSP kept at 4°C, were warmed to room temperature. Rat pups were maintained at warm (37°C) in a parallel transparent box next to the mother box during one hour for gastric emptying. Mixtures of miRNAs and DOSP were extemporally prepared. We have chosen an oral administration at the beginning of the dark phase.^20^ Gastric fluids were collected to evaluate a putative re-export of loaded miRNA though extracellular vesicles. In the gut, we have sampled the stomach as the inoculation site and the duodenum as relevant for our transgene expression. The rationale for choosing brain area was for the hypothalamus as the main center of energy homeostasis, the hippocampus as involved in the memory of food reward and choice, the brain stem as the outcome of the vague nerve linking the intestine to the brain. We have screened the effects according to gene sets related to inflammatory and enteroendocrine status of stomach or duodenum, and on the serotoninergic/dopaminergic balance of brain areas. Our transgenic GFP-CCK-p rat derived from the Sprague-Dawley strain is allowing to follow an enteroendocrine cell lineage labeled with Green Fluorescent Protein in duodenum crypts. In addition, the Green Fluorescent Protein transcripts can be followed in rat neuronal cells expressing CholeCystoKinin (CCK). B. Transgenic rat generation. Targeted integration of a CCK promoter-GFP cassette into the rat Rosa26 locus (left panel). Schematic representation of the rat Rosa26 locus. TALEN cleavage (vertical arrow) in the first intron, as well as the sequences recognized by both TALENs, the targeting vector with expression cassette (3878 bp) and the 5′ and 3′ homology arms (HA) (800 bp each) are indicated. The PCR primers flanking the cleavage sequence used to do the first genotyping are also indicated (right panel). Schematic representation of the CCK-GFP cassette integration. To verify the integrity of the CCK-GFP cassette, genomic DNAs were PCR amplified with the primers described and the PCR amplicons were analyzed for their size and Sanger sequences.

As shown in Figure 1B, a Sprague-Dawley transgenic strain of rats expressing enhanced green fluorescent protein eGFP under the control of the CCK promotor, generated using the transcription activator-like effector nuclease (TALEN) methodology and knock-in in the *Rosa26* locus^38^ was selected. This transgenic rat line allows studying CCK-eGFP+ cells, for example in duodenum crypts and villi.^39^ In addition, the eGFP (*gfp*) transcripts can be followed in rat neuronal cells expressing CCK.^40^ PCR for genotyping and qPCR primers are shown in Supplementary Table 1.

### 2. miRNAs

Two miRNAs, present in breast milk, were used for oral supplementation, miR-320-3p (MIMAT0000903 AAAAGCUGGGUUGAGAGGGCGA) already described by us and others as having epigenetic activity^20, 24^, and miR-375-3p MIMAT0005307 UUUGUUCGUUCGGCUCGCGUGA, known to target proliferative cells in gut and related to Vitamin-E metabolism in humans.^41^ The rno-miR-320-3p or rno-miR-375-3p were ordered from Eurofinns, Germany and they were checked at reception by reverse transcription and q-PCR with corresponding TaqMan probes (Supplementary Table 1).

### 3. Ribonucleic acid vector

The vector was Di-Oleyl-Succinyl-Paromomycin (DOSP), used for *in vivo* short-term transfection of miRNA^20^ and of mRNA.^18^ The vector is non-cytotoxic and allowed in food practice. Prior to use, the quality of the vector was assessed by the size distribution of DOSP nanoparticles with a pike at 200 nm on a Gold-q-Nano (Izon).

### 4. Global analysis of miR-320-3p or miR-375-3p networks

#### 4-1. miR Target enrichment analysis

To identify potential target genes regulated by miR-320-3p or miR-375-3p, the miRNA-target pairs were retrieved from TargetScan 7.2 (up date: march 2018) (http://www.targetscan.org), miRWalk v6.0, (update: Jan 2021 www.umm.uni-heidelberg.de/apps/zmf/mirwalk), and miRDB v6.0 (up date: June 2019, http://mirdb.org/), exploring the miRNA binding sites within the complete sequence of rat genome (including 5′-UTR, 3′-UTR regions as well as coding sequences) and combining this information with a comparative analysis of predicted binding sites. The three databases were jointly mined, and overlaps of the results were generated to obtain the list of the most potential regulated transcripts.

#### 4-2. Pathway enrichment analysis

To identify pathways in the list of the potentially regulated mRNA we used Panther v 16.0 (released 2020-12-01, http://www.pantherdb.org/). To ensure the validity of our findings, we only considered the three pathways more relevant to both miRNAs.

### 5. Gastric fluids and Tissue samples

Briefly at sacrifice, the stomach content was collected and stored under liquid nitrogen.^20^ Before analysis, the samples were thawned and extracellular vesicles were recovered by elution through a qEV column (Izon) and processed for analyzing the size distribution with a qNano (Izon). The stomach and the duodenum were rinsed with a Phosphate-Buffered saline solution free of calcium and magnesium ions. Pieces of the lower part of the stomach (fundus) and of the duodenum were immersed under liquid nitrogen. In addition, pieces of the duodenum were fixed in 0.1 M PBS containing 4% paraformaldehyde for 24h and then embedded in paraffin for histology. The incidence of miRNA/DOSP bolus was evaluated 8 hours after inoculation by measuring D-Glucose in blood, the contents of miR-320-3p and 375-3p in exosome fractions of gastric fluids, and in rat pup plasma. For rats on Day-45, in addition to stomach and duodenum, brain compartments were immersed directly under liquid nitrogen.

### 6. Analysis of miRNA and mRNA by q-PCR

Total RNA extraction was done with Qiazol (Qiagene, France), and cDNA was obtained with TaqMan miRNA kit (Thermofisher, France). All primers are listed in Supplementary Table-1. We have used either TaqMan’primers, or self-designed primers with SyberGreen. At the taxonomic level, the miR-320-3p is a non-canonical miRNA^42^ with a non-described 5p form in humans and rats. In the bioprocessing of true miRNAs like miR-375, both the 3p and 5p molecules are expressed allowing quantitative exploration by the northern blot of their ratio in human cell lines.^43^

### 7. Chromatin ImmunoPrecipitation

The methylation of the Histone 3 in the lysine residue 4, generally, is associated with the activation of transcription of nearby genes and we used Pierce kit methodology.^20^ It should be underlined that little is known concerning the nuclear delivery of the non-complexed miRNA.^46, 47^ Chromatin immunoprecipitation was performed with Pierce Chromatin Prep Module (Thermo Scientific #26158). Briefly, small tissue aliquots were cross-linked by exposition to 1% formaldehyde.

Chromatin was fragmented by Micrococcus Nuclease.

Immunoprecipitations were performed using 1 μg of Anti-Trimethyl-Histone-3-Lys-4 (-Thermo Fisher Scientific; catalog# PA5-17420) overnight at 4 ◦ C. Micrococcus nuclease was used at 0.25 μL/sample.

We collected immune complexes with agarose A/G for 2 h at 4 ◦ C, beads were rinsed twice by PBS0 and pelleted at 94 × g for 1 min. Immune complexes were eluted by adding 100 μL of elution buffer to pelleted beads. After brief vortexing, preparations were incubated at Room Temperature for 15 min.

Thereafter, beads were spun down and the supernatants (eluates) carefully transferred to another tube. The elution step was repeated. Both eluates were combined.

We added 5 M NaCl and proteinase K allowing crosslink reversion by 1.5-h incubation at 65 ◦ C. Nucleic acids were recovered by Qiagene miRNA-Easy kit and analyze ChIPped chromatin using quantitative PCR. iQ SYBR Green Supermix (Bio-Rad) was used to perform real-time PCR on an iCycler iQ system (Bio-Rad) with promoter-specific primers (Supplementary Table-1).

### 8. Density of duodenal green-labeled CCK-p enteroendocrine cells

Following rehydration, thick sections (4 um) were stained with chromogranin A (marker of total enteroendocrine cells) rabbit polyclonal antibody (diluted 1/500; 20085 Immunostar). After incubation with a secondary biotinylated goat anti-rabbit (diluted 1/1000, A24541, Life Technologies), chromogranin A was revealed with Alexa Fluor 568-conjugated streptavidin (S11226, Invitrogen). Slides were mounted in Prolong Gold antifade reagent (Invitrogen) that contains DAPI to counterstain nuclei. The density of CCK-producing cells that stained in green (endogenous GFP) and positive for chromogranin A was measured by fluorescence microscopy (Zeiss, Axio Imager M2m) in the crypts of 3 sections of duodenum using a 40 X objective. The data were expressed as the percentage of chromogranin A-positive cells or CCK (GFP) positive cells per crypt.

### 9. Sample nomenclature, selection of reference genes, statistical analysis

The nomenclature for identifying sample is: stomach “sto”; brain stem “bs”; hippocampus “hip”; hypothalamus “hy”; weaning at Day-15 “w15”; weaning at Day-30 “w30”; oral bolus with miR-320-3p/DOSP “b320”; oral bolus with miR-375-3p/DOSP “b375”; oral bolus of controls “btem”. As an example, hy-w15-btem means that the rat sample is from Hypothalamus, from a rat weaned at Day-15 receiving a control bolus. We are using 3 miRNAs as reference genes.^20^ But the miR-146b-5p has been reported as a marker of depression.^48^ So, we have also analyzed the datasets using either all miRNAs or all mRNAs per sample as a global normalization procedure. By applying this technique with miR-146b-5p, we did not find significant variation. Consequently, the miR-146b-5p remains under our hand, a valid reference gene (unshown results). We have decided to analyze Delta-Cq after a log10 transformation^20^, justifying our use of one way ANOVA for the comparison between groups, and to test multiple comparisons. Data analysis was performed with R Commander and R suite or Graph Pad software.

We have used Cytoscape to create networks of miRNA, mRNA significantly deregulated in our data-set. Excel files of raw Cq data organized by tissues can be made be accessible at the UN-Cloud of the Nantes University.

## Results

### 1. In silico analysis of rno-miR-320-3p and rno-miR-375-3p networks

We have found that the sequence coding for mature miR-320-3p was identical in rat, mouse, and human. Pairwise alignment revealed that miR-320-3p is antisense, encoded in the intergenic region of chromosome 8 at approximately 200 bp upstream of the TSS of RNA polymerase III subunit D (polr3d) in the chromosome 15 in rat (rn6), the human (CRCh38.p13), and the chromosome 14 in mouse.^24^ We have found that the sequence coding for mature miR-375-3p was identical in rat mouse, and human. The mature full sequence of miR-375-3p was located at chromosome 9 for rat, at chromosome 2 for humans, and chromosome 1 for mice. In the three genomes, it is located at the non-coding region.

Current miRs target prediction algorithms regularly present different numbers of potential interactions. Due to that, we have fused the results of three databases to obtain the most accurate list of target genes. For miR-320-3p, the list consisted of 111 transcripts distributed between 69 pathways. For miR-375-3p, we obtained a list of 24 transcripts found in 12 pathways.

By target mining miRWalk on 3’-UTR, set on perfect matching between miRNA and target mRNA, we have found 126 genes common between miR-320-3p and miR-375-3p. They are corresponding to the Kegg pathway : rno01100_Metabolic_pathways.

We have found 308 genes common between, miR-320-3p and miR-132-3p, among which we are exploring PER2. We have found 112 genes common between miR-320-3p and miR-16-5p, among which we are exploring period2 (per2), circadian locomotor output cycles kaput (clock).

The miRNA delivered to cells with appropriate carriers or expressed in cells using suitable vectors often triggers both intended sequence-specific silencing effects and unintended sequence-non-specific immune responses.^44^ So we have established a list of genes for exploring the inflammatory status of stomach samples: Interleukin1A (*IL1A*), Interleukin6 (*IL6*), Interferon-gamma (*IFNg*), Signal transducer and activator of transcription 3 (*stat3*), Interleukin10 (*IL10*), Tumor Necrosis Factor alpha (*tnf-a*), Signal transducer and activator of transcription 1 (*stat1*), *iNOS, PPARg* (peptide related to food consumption), *foxa1*, Interleukin1B (*IL1B*). By data mining, these genes are related to Inflammatory status: GO:0006954_inflammatory_response and GO:0005125_cytokine_activity.

In brain samples, we have explored *clock* gene which is common to miR-320-3p and miR-375-3p, along with Brain And Muscle ARNT-Like 1 (*bmal1*), Period1 (*per1*), and Period2 (*per2*). Interactions between miRNA and mRNA were built using miRWalk^45^, and for the serotoninergic/dopaminergic profiles: serotonin transporter (*sert*), 5-hydroxytryptamine receptor 1B (*5ht1b*), 5-hydroxytryptamine receptor 2C (*5htr2c*), Dopamine receptor D1 (*drd1*), Dopamine receptor D2 (*drd2*), Cholecystokinin (*cck*). They are found by data mining related to GO:0007420_brain_development; GO:0003676_nucleic_acid_binding, and GO:0006357_regulation_of_transcription_by_RNA_polymerase_II.

Taking advantage of our transgenic rat, we have assayed markers of enteroendocrine lineage (Paired box gene 4 *(pax4*), Paired box gene 6 (*pax6*), ghrelin (*ghrl*), Peptide YY (*pyy*), chromogranin A (*chgA*), Gastric Inhibitory Polypeptide (*gip*), Cholecystokinin (*cck*) by q-PCR.

In summary, our *in silico* analysis is showing that miR-320-3p is influencing on average a gene network four times wider than miR-375-3p.

### 2. Immediate effects of force-feeding miRNAs/DOSP in the stomach and duodenum of breast-fed rats at Day-12

The transgenic rat strain was checked for correct expression of the transgene by PCR using tail biopsies to check for the homozygote status. Duodenal cross-section and immunostaining were realized on D12 and D45 rats to check for the expression of GFP-labelled duodenal cells according to the co-expression of *chgA*, a biomarker of total enteroendocrine cells (unshown results). As shown in Supplementary Figure 1, the relative level of miR-320-3p is increased in the stomach wall of rat pups supplemented with miR-320-3p/DOSP (B, p=0.05) whereas hspb6 transcripts are decreased (D, p<0.05) with a similar trend for polr3d mRNA (C). Concerning enteroendocrine markers, only *chgA* was highly up-regulated (p<0.0001) with miR-320-3P/DOSP treatment. Likewise, the *IL1B* was down-regulated only with rat pups treated with miR-320-3P/DOSP (p<0.001).

Moreover, the treatment with miR-320-3p/DOSP induced a significant decrease in chromatin complexes harboring H3K4me3 and polr3d promoter in gastric cells (Supplementary Figure 1E, p<0.05). Surprisingly, we did not detect any immediate effect of miR-375-3p/DOSP both on transcripts or on chromatin complexes (Supplementary figure 1E), but we infer from the long-term effects reported below that the delivery of miR-375-3p was done according to its putative cytoplasmic site of bioactivity. We did not detect leakage of both miRNAs in exosomal fractions of gastric fluids, confirming our previous demonstration that the vector delivered the miRNAs into the cytoplasm of digestive cells.^20^ The levels of D-Glucose were not different (Group average ± standard deviation, treated with miR-320-3p: 135 mg D-Glucose/dL±10,9; treated with miR-375: 131,2 ± 15; control: 127,2 ± 6,5). The levels of miR-320-3p or miR-375-3p in plasma according to treatment with miR-320-3p/DOSP (Average Cq ± standard error: 22.79 ± 0.72or 29,57 ± 0.65), miR-375-3p/DOSP (21.99 ± 0.89 or 27.78 ± 1.34), or control (21.83 ± 1.04 or 28.38 ± 0.84) were not different.

In summary, these data on the immediate effect of rno-miR-320-3p/DOSP suggest that the miRNA molecules are bioactive both in the cytoplasm and chromatin complexes, but we did not show any evidence of an immediate effect of rno-miR-375-3p/DOSP at the transcript level in the stomach, even on enteroendocrine markers.

### 3. Effects of early weaning in control groups (not supplemented with miRNA)

#### 3.1 Stomach

Gastric endogenous miR-320-3p and miR-375-3p were not significatively different between early-weaned and regularly weaned rats at Day-45 (Figure 2A; note that the miR-132-3p or miR-504 were not detected in stomach samples). No difference between weaning times has been recorded for the expression of *polr3d* or *hspb6* mRNA (Supplementary Figure 2A, B). The level of *tnf-a* was down-regulated, and those of *IL6*, and *IFN-g* were up-regulated in early-weaned controls compared to the regular weaning controls (Figure 3A; Supplementary Figure 2). The altered expressions of these cytokine transcripts suggest a long-lasting state of inflammation induced by early weaning stress. The relative levels of chromatin complexes harboring H3K4me3 were slightly lower in the early-weaned rats (Supplementary Figure 1F, G).

**Figure 2.**
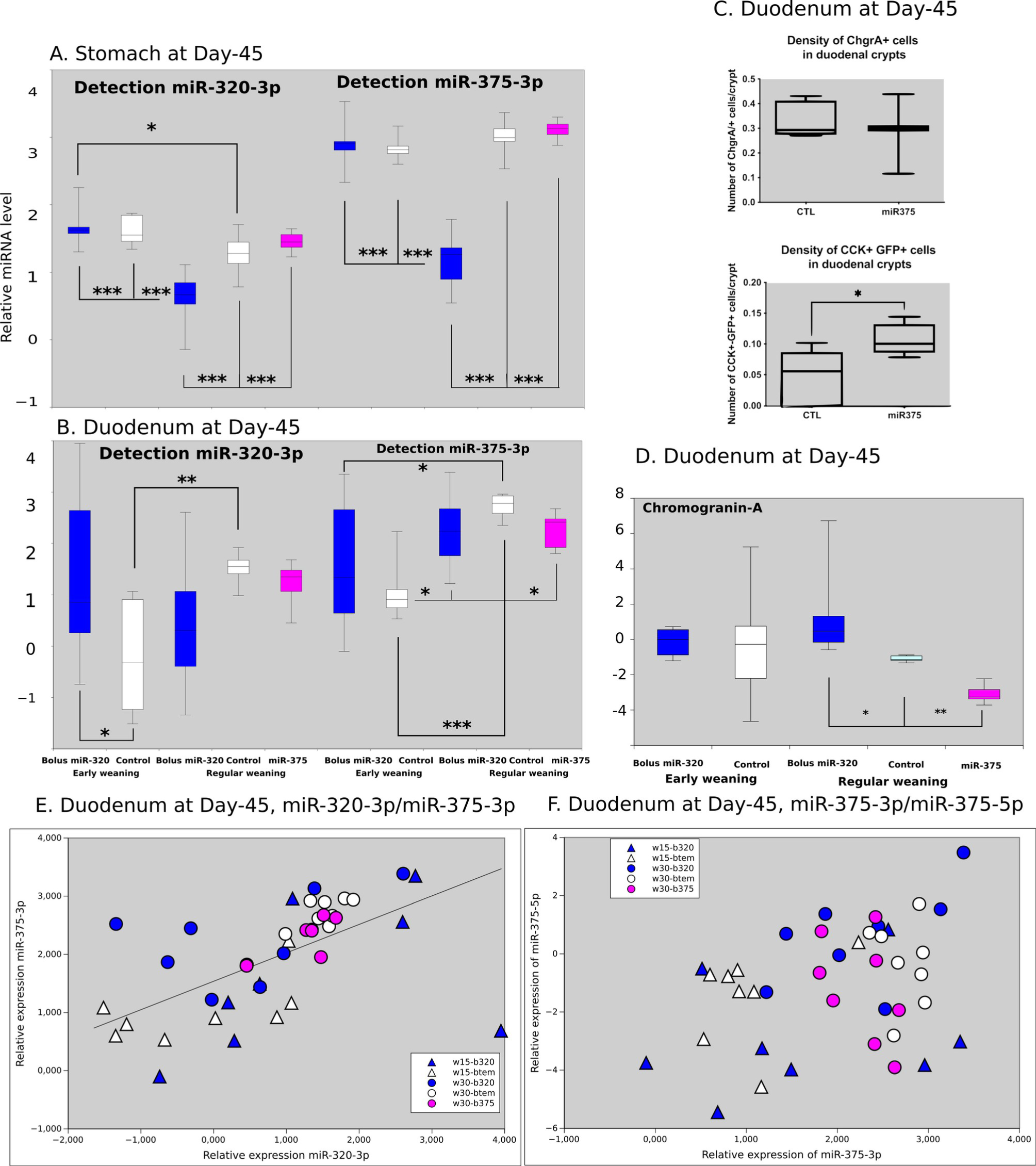
Long term effects of miR-320-3p/DOSP or miR-375-3p/DOSP on miR-375-3p and miR-320-3p expressions in stomach (A), duodenum (B), long term effect of miR-375-3p/DOSP followed by regular weaning on the density of CCK+GFP+ duodenal cells (C) or long term effect of miR-320-3p/DOSP or miR-375-3p/DOSP followed by early or regular weaning on the level of chromogranin A transcripts in duodenum (D). Scatter plots between miR-320-3p/miR-375-3p and miR-375-3p/miR-375-5p are shown in E and F, respectively. In E, low correlation (R2=0.54, black line) and in F, the miR-375-5p levels of w15-b320 were down-regulated in comparison with w30-b320 (p=0.02), and w30-btem (p=0.04). The light gray background reminds that rats were sacrificed in the dark phase. Note: * p<0.05; ** p<0.01; *** p<0.001.

**Figure 3.**
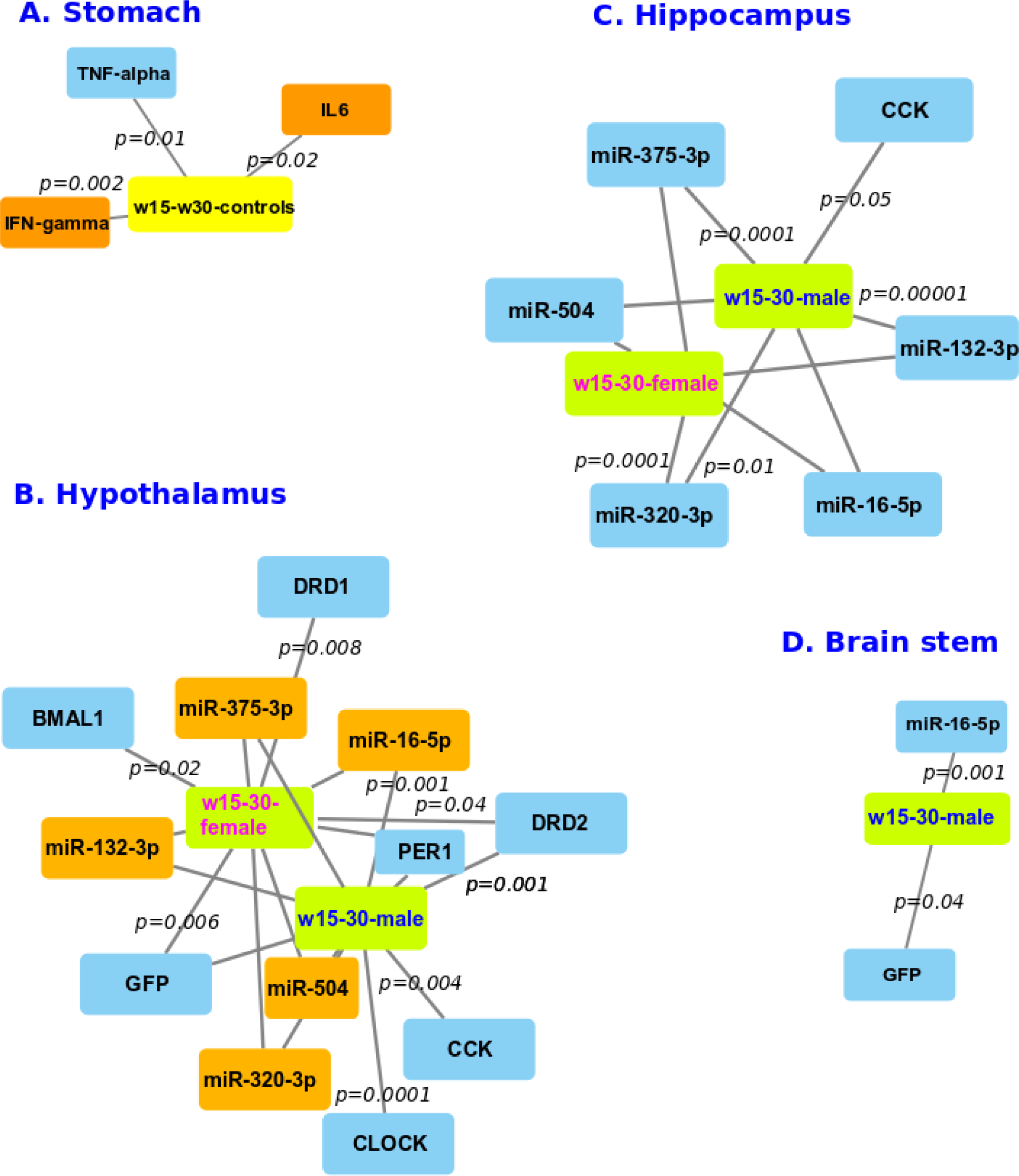
Comparison of early and regular weaning controls. Network of genes significantly deregulated in stomach wall (A), hypothalamus (B), hippocampus (C), and brain stem (D). Note on group nomenclature, for instance, “w15-w30” meaning comparison between the early weaned controls with the regular weaning controls. Edge length is inversely proportional to p significant threshold. Most p values are indicated in italic. Orange up-regulated, blue down-regulated.

#### 3. 2. Duodenum

Endogenous miR-320-3p and miR-375-3p at Day-45 (Figure 2B, p<0.05 and p<0.001, respectively) were down-regulated in early weaned rats as compared to regularly weaned ones. Consequently, these miRNAs could be crucial in the peripheral response to Early Life stress. We did not find strong correlation between the relative levels of miR-320-3p and miR-375-3p (Figure 2E) nor between miR-375-3p and 5p (Figure 2F). But the miR-375-5p level of w15-b320 were down-regulated in comparison with w30-b320 (p=0.02), and w30-btem (p=0.04, Figure 2F).

#### 3. 3. Brain

A comparison of controls submitted to early or regular weaning showed drastic differences between hypothalamus, hippocampus, and brain stem.

In the hypothalamus, all tested miRNAs (miR-320-3p, miR-375-3p, miR-16-5p, miR-132-3p, and miR-540 were up regulated in both females and males (Figure 3B, 4A, 5A). We have found a down regulation of *drd1* in the hypothalamus of early-weaned females with a higher expression of miR-504 according to Huang and Li, 2009^4^ (Figure 6A, the correlation between *drd1* and miR-504 for 39 animals was of -0.755 irrespectively of their experimental group). However, we did not find any difference in *drd1* relative level in the male hypothalamus according to weaning time, even if an up-regulation of miR-504 was recorded (Figures 3B, 5A, Supplementary Figure 3A, B). If all tested miRNAs (miR-504, miR-16-5p, miR-132-3p, miR-320-3p, miR-375-3p) were all up-regulated in the hypothalamus of early-weaned rats at Day-45 of life, only the *per1* transcripts of the circadian clock were down-regulated for male and female rats (Figure 3B; Supplementary Figure 4). However, the females had a down-regulation of *bmal1*, and the males of *clock* (Figure 3B; Supplementary Figure 4). The *cck* transcripts were down-regulated with early-weaned hypothalamus, and according to the logic of the transgene construct, a down-regulation of the *gfp* transcripts was also found in males and females. No strong correlation between *cck* and *gfp* (correlation of 0.537) was recorded, suggesting that if the promoter was driven by the same transcriptional machinery as the *cck* gene, the transgene was independently regulated from the *cck* endogenous gene promoter (Supplementary Figure 5).

**Figure 4.**
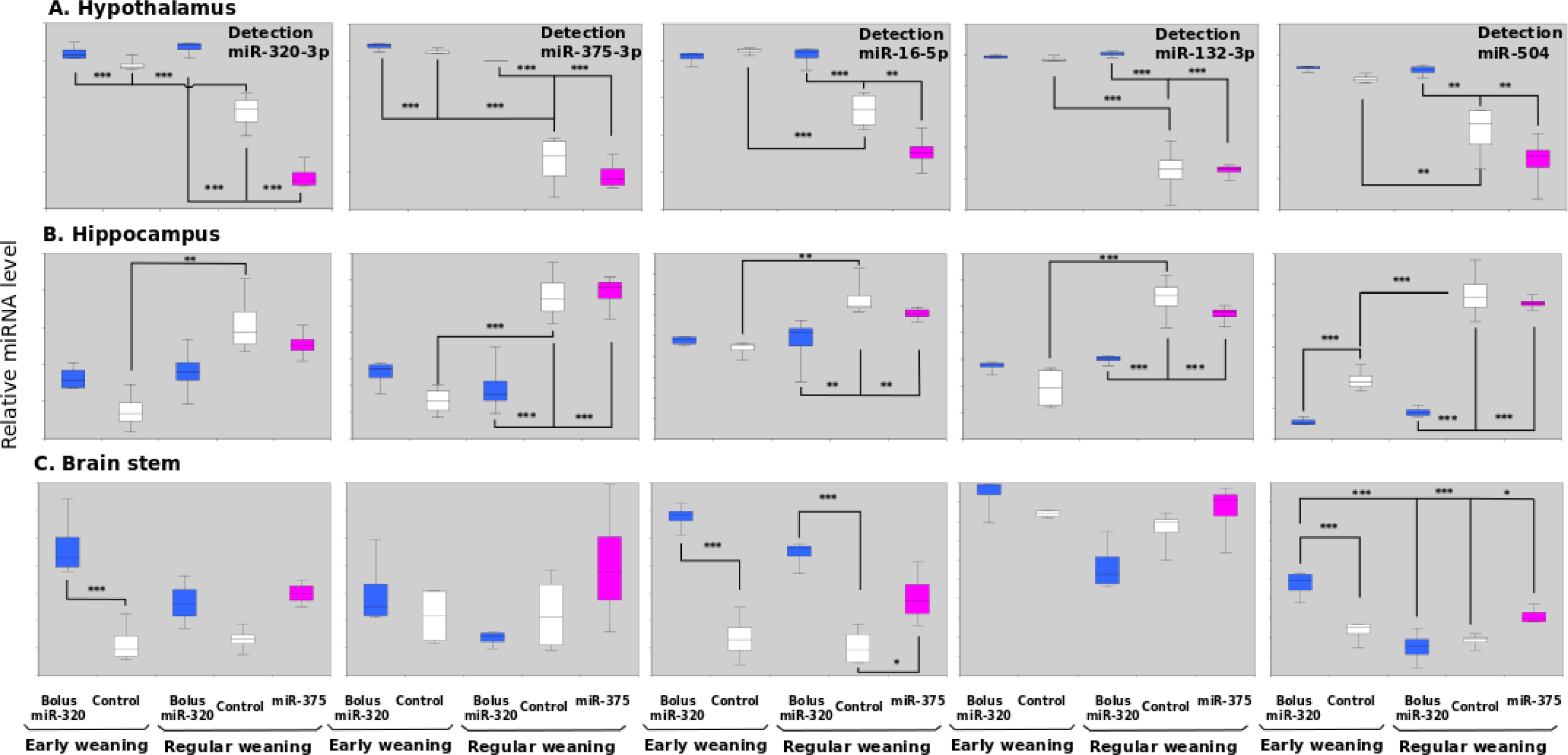
Evolution of miRNAs in hypothalamus (A), in hippocampus (B), and in brain stem (C) of females treated with miR-320-3p or 375-3p/DOSP according to early or regular weaning. The light gray background reminds that rats were sacrificed in the dark phase. Note: * p<0.05; ** p<0.01; *** p<0.001.

**Figure 5.**
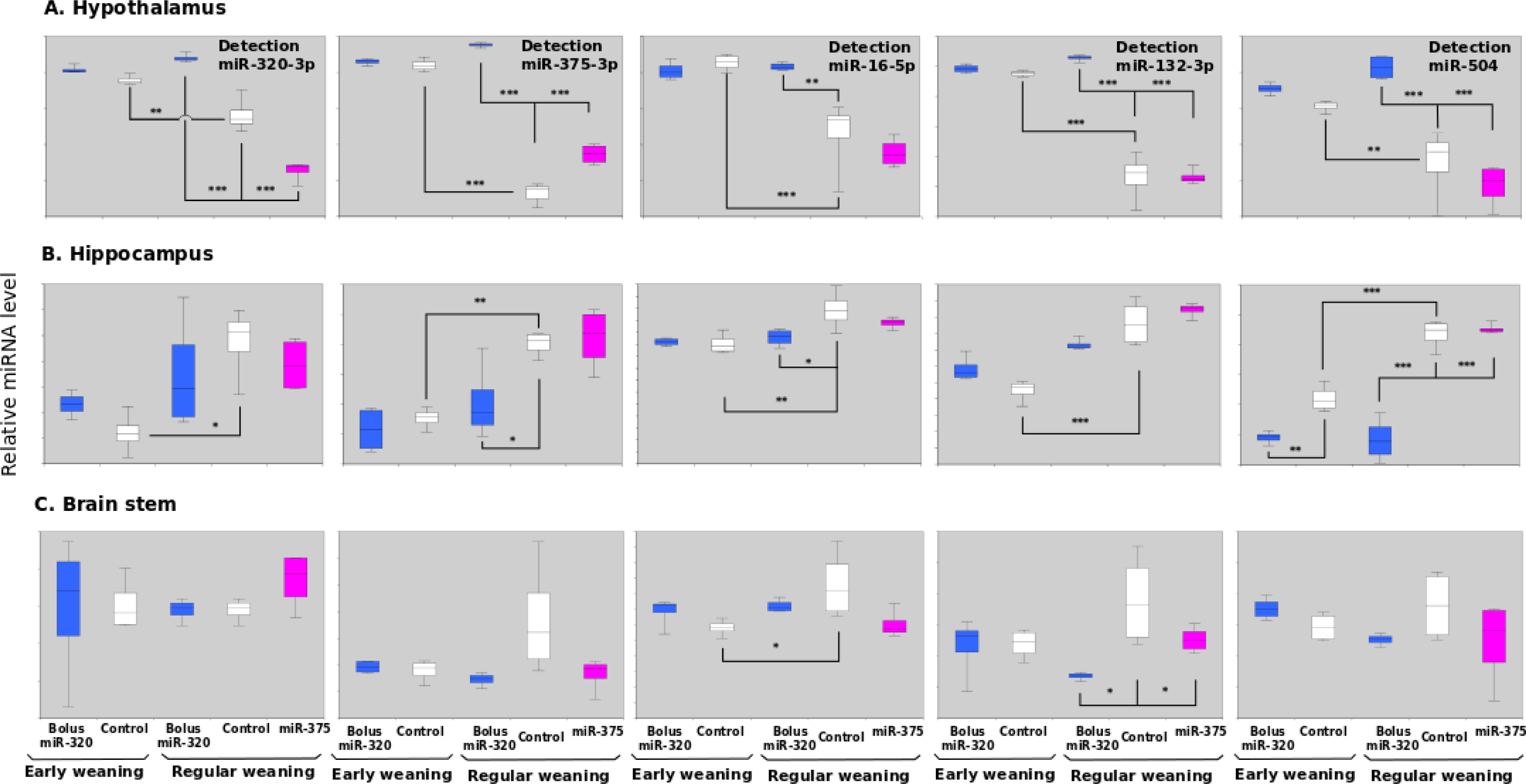
Evolution of miRNAs in hypothalamus (A), in hippocampus (B), and in brain stem (C) of males treated with miR-320-3p or 375-3p/DOSP according to early or regular weaning. The light gray background reminds that rats were sacrificed at ZT-20H in the dark phase. Note: * p<0.05; ** p<0.01; *** p<0.001.

**Figure 6.**
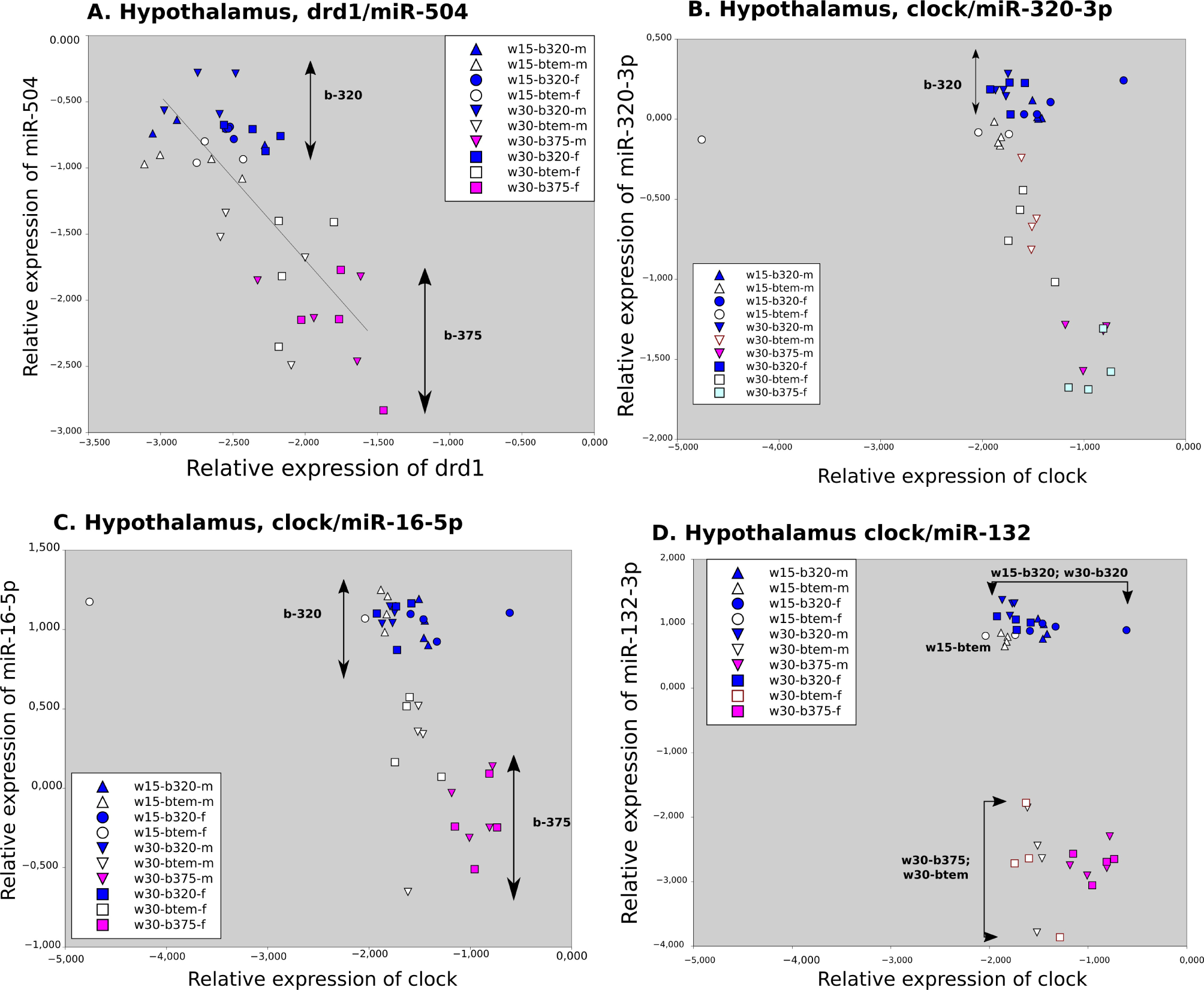
Long term effect of miR-320-3p/DOSP according to early or regular weaning. Note the negative correlation (R=-0.75, linear regression in black) between miR-504 and DRD1 transcripts (A), and the up-regulation of miR-320-3p (B), miR-16-5p [C] and 132-3p (D) for early weaned rats and regularly weaned rats treated by miR-320-3p/DOSP in hypothalamus cell extracts. The level of clock transcript is significantly different only in B and C. The light gray background reminds that rats were sacrificed in the dark phase. Note: * p<0.05; ** p<0.01; *** p<0.001.

In the hippocampus, all miRNAs were down regulated (Figure 3C, 4B, 5B) like for miR-320-3p and miR-375-3p in the stomach. In Figure 3C, all tested miRNAs were down-regulated for both sexes, as well as for *cck* gene (Supplementary Figure 5).

In the brain stem, only miR-16-5p in males was down regulated (Figure 3D; 4C). In Figure 3D, only miR-16-5p and *gfp* were down-regulated. These results obtained at Day-45 for Sprague-Dawley rats identify new molecular pathways in the follow-up of our work^37^, realized on Wistar rats between Day-250 and 300.

In summary, after early weaning, endogenous miR-320-3p and miR-375-3p are deeply altered in the duodenum, hypothalamus, and hippocampus of young rats. These data are strongly supportive of using these milk miRNAs as supplements in lactating rat pups, even with discordant results on stomach (non-significative effects for miR-320-3p or miR-375-3p in Figure 2A) and duodenum (significative effets for miR-320-3p in Figure 2B).

### 4. Comparative evaluation of the long-term effects induced by miR-320-3p/DOSP or miR-375-3p/DOSP, with subsequent early or regular weaning

#### 4-1. Evaluation of transcripts on stomach or duodenum cell extracts and of duodenal cell density by immunhistochemistry and expression of duodenal GFP transcripts

In the stomach, miR-320-3p and miR-375-3 transcripts were significantly down-regulated in young rats when treated with miR-320-3p/DOSP compared to controls and miR-375-3p/DOSP (Figure 2A, p<0.001). We did not observe in young rats, deregulation of miR-375-3p, nor of miR-320-3p with rat pups forced-fed with miR-375-3p/DOSP. By contrast, the miR-320-3p/DOSP treatment had significantly up regulated the endogenous miR-320-3p transcripts in the duodenum of early-weaned rats (Figure 2B, p<0.05).

Moreover, as shown in Figure 7A, the enteroendocrine markers (*pax4*, *pax6*, *chgA*) and *tnf-a* were all down regulated after treatment with miR-375-3p. By contrast, the treatment by miR-320-3p/DOSP down-regulated miR-320-3p (p=0.001), up-regulated in parallel miR-375-3p (p=0.00001). The *tnf-a* was up-regulated (p<0.0001) along with a down-regulation of *stat1* (p=0.05), of *IL10* (p=0.00001), and of *foxa1* (p<0.001) for rats treated with miR-320-3p/DOSP and weaned at Day-30 (Supplementary Figure 2C, D). For early-weaned rats treated with miR-320p/DOSP, these molecules were not significantly altered, suggesting that the inflammatory status was unchanged by the miRNA supplementation. The *grlh* and *pyy* transcripts were down-regulated respectively at p=0.04 and p=0.03. We have found a trend to a down-regulation for the GFP-CCK-promoter transcripts and a strong correlation between *cck* and *gfp* (correlation of 0.937 at Day-12) without any strong difference for the correlation (0.91 at Day-45) after miR-375-3p/DOSP or controls submitted to regular weaning.

**Figure 7.**
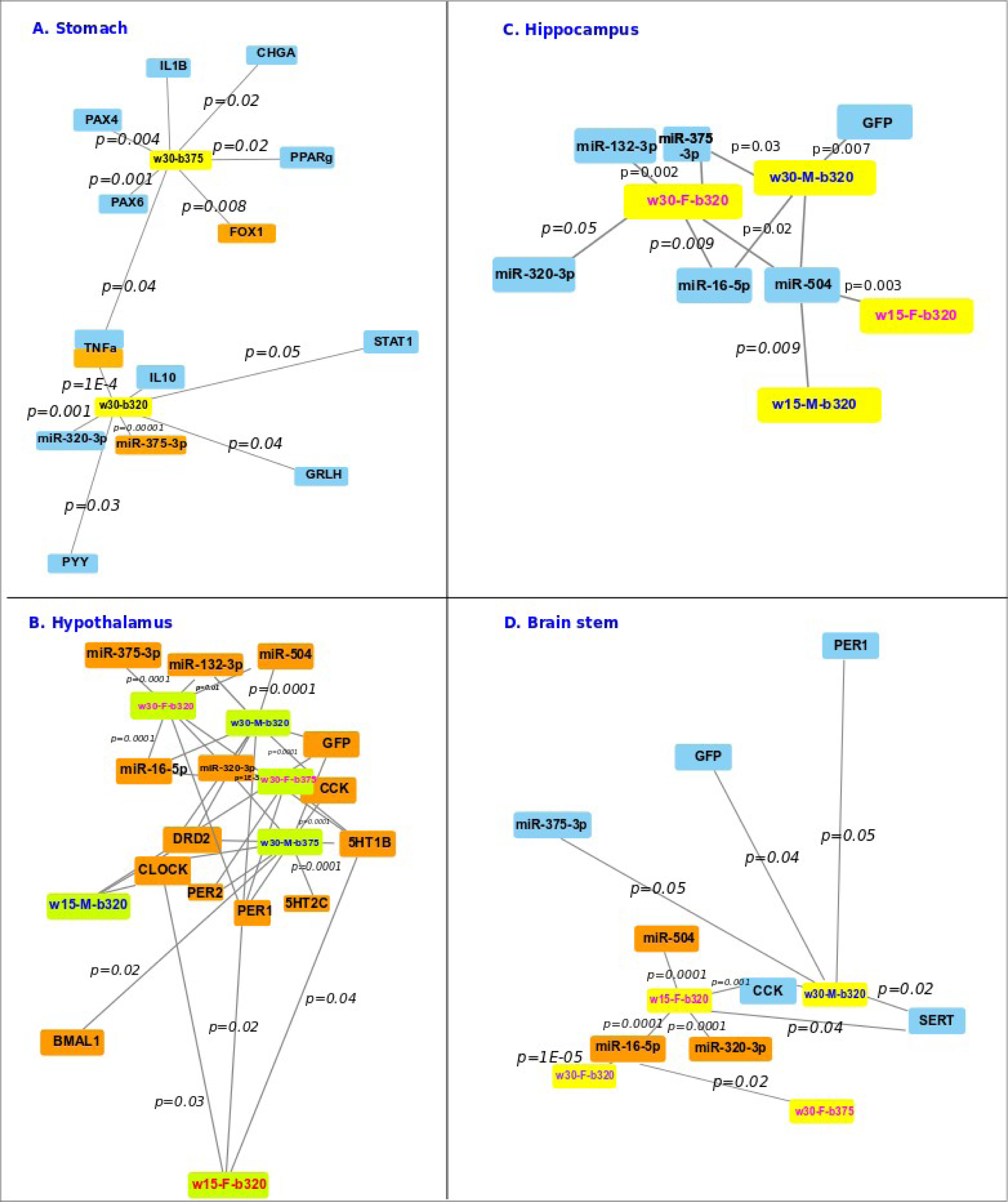
Long term effect of miR-320-3p/DOSP according to early or regular weaning and of miR-375-3p/DOSP with regular weaning. Network of genes significantly deregulated after treatment with miR-320-3p/DOSP or miR-375-3p/DOSP comparatively to control in the stomach wall (A), hypothalamus (B), hippocampus [C], and brain stem (D). Note on group nomenclature, for instance, “w15-w30” meaning comparison between the early weaned controls with the regular weaning controls. Edge length is inversely proportional to p significant threshold. Most p values are indicated in italic. Orange up-regulated, blue down-regulated.

We did not detect any difference between chromatin complexes harboring H3K4me3 and polr3d promoter in gastric cells after oral administration of synthetic miR-320-3p nor of miR-375-3p and according to the weaning periods (Supplementary Figure 1F, G). After regular weaning, no difference between polr3d, nor hspb6 transcripts at Day-45 was found both in the stomach and the hypothalamus.

In the duodenum, a long-term effect of miR-375-3p/DOSP was noted on the density of GFP-CCK-p labeled duodenal cells (Figure 2C, p<0.05). As shown in Figure 2D, the miR-320-3p/DOSP treatment increased the level of *chgA* and the miR-375-3p/DOSP treatment decreased both Chromogranin A (Figure 2D, p<0.01) and Gastric Inhibitory Polypeptide (p<0.001).

In summary, long-term effects of an oral exposure during lactation to the high concentration of miR-320-3p were found with unforeseen consequences on the transcript levels of endogenous miR-320-3p and miR-375-3p on the group with regular weaning. The most striking result is about the up regulation of miR-320-3p in the duodenum suggesting that the stem cell compartments along the gut were differently altered by the treatment with miR-320-3p/DOSP.

#### 4-2. Evaluation of transcripts on cell extracts of the hypothalamus, hippocampus, and brain stem

In the hypothalamus of females (Figure 4A) and males (Figure 5A) treated with miR-320-3p/DOSP, the endogenous miR-320-3p transcripts were up-regulated like miR-375-3p, miR-16-5p, miR-132-3p, and miR-504 (all p < 0.001; correlation coefficients between all the miRNAs were positive and superior to 0.88). We did not detect any difference for miR-320-3p transcripts for both sexes with the groups supplemented with miR-375-3p/DOSP. By contrast, we had a strong down regulation of miR-375-3p for the males or the females treated with miR-375-3p/DOSP.

Interestingly, the miR-375-3p levels were altered for both sexes when treated with miR-320-3p/DOSP and weaned at Day-30. The males treated with miR-320-3p/DOSP and miR-375-3p/DOSP had an altered level of miR-320-3p, but the observation was true only for the females treated with miR-320-3p/DOSP. Surprisingly, we did not observe any strong effect of miR-320-3p/DOSP supplementation on young rat hypothalamus enduring early-weaning, as the level of endogenous miR-320-3p was already very high (Figures 4A, 5A, 6A).

All males had a deregulation of clock and drd2 transcripts when supplemented with miR-320-3p/DOSP and, for the ones with regular weaning, with miR-375-3p/DOSP supplementation (Figure 7B, Supplementary Figures 3, 4). Unlike miR-504 with *drd1*, the levels of miR-320-3p, miR-16-5p or miR-132-3p were not correlated with the level of *clock* transcripts (Figure 6B-D). The females having endured both stress (miR-320-3p/DOSP and early weaning), displayed alterations in 5HT1B level like females with regular weaning and in PER1, another gene of the circadian clock. Males and females treated with miR-375-3p/DOSP displayed alteration in *per2* (Figure 7B, Supplementary Figure 4).

In the hippocampus, the levels of miR-320-3, miR-375-3p, miR-16-5p, miR-132-3p, and miR-504 transcripts were all down regulated for the group treated with miR-320-3p/DOSP (Figures 4B; 5B; all p at least below 0.01). By contrast, the group treated with miR-375-3p/DOSP were for all miRNAs in the range of controls. The miR-504 and the miR-132-3p were deregulated for male and female weaned at D-30 treated with miR-320-3p/DOSP, but the rats weaned at Day-15 did not shown any difference for miR-504 and miR-132 (Figures 4B; 5B; 7C).

In the brain stem of females, a significant up regulation was observed with miR-320-3p for the females treated with miR-375-3P/DOSP. An up regulation of miR-16-5p and miR-504 with the females treated by miR-320-3p/DOSP. With males, a significant down regulation of miR-132-3p transcripts was noted with the group treated by miR-320-3p/DOSP. Both male and female early-weaned rats displayed a deregulation of *cck* and *sert* (Figure 7D, Supplementary Figure 5). All females had a down-expression of miR-16-5p when supplemented with miR-320-3p/DOSP and, for the ones treated by miR-375-3p/DOSP with regular weaning. The *gfp* transcripts were significantly down regulated for the males supplemented with miR-320-3p/DOSP and after regular weaning (p<0.05). On Figure 7D representing hippocampus data, the miR-504 was down regulated for all groups when supplemented with miR-320-3p/DOSP indicating a strong effect of this miRNA supplementation. It should be underlined that no difference was found between sex and weaning times.

Our transgenic rat express the *gfp* transcripts in all cells permissive for CCK promoters beside the enteroendocrine lineage of duodenum. As such, we have found a deregulation for all male young rats after early or regular weaning, treated with miR-320-3p/DOSP and miR-375-3p/DOSP, but a similar deregulation was circonscribed to the females treated with miR-320-3p and early-weaned. It should be underlined that *gfp* was altered both in hippocampus and brain stem with males supplemented with miR-320-3p/DOSP.

With early weaning, the levels of miR-320-3p were up regulated in duodenum extracts of miR-320-3p/DOSP treated rats (Figure 2B). However, we did not find any difference between miR-320-3p/DOSP and controls in stomach extracts (Figure 2A), as well as for miR-16-5p (Mean Cq ± sme of w15-b320 (19.83±1.65) against w15-btem (20.16±2.01)) nor in the duodenum extracts (unshown results). Likewise, in the hypothalamus of both sexes (Figures 4A, 5A) or with hippocampus, except for a down regulation of miR-504 for both sexes (Figures 4B, 5B). In the brain stem of females, a significant up regulation was found for miR-320-3p, miR-16-5p, and miR-504. A down regulation was found for the relative expression of *cck* and *sert* transcripts. A trend of up regulation was observed with miR-375-3p and miR-132-3p (Figures 4C, 5C). The significant evolution of gene expression is summarized Figure 7. In addition, we did not find any difference for sidt1 between early weaned rats treated with miR-320-3p/DOSP and corresponding controls (a trend of down-regulation was found with sidt1: Average Cq ± sme of w15-b320 (32.55 ± 4.59) against w15-btem (26.19 ± 7.54). We have assayed miR-375-5p finding a huge variability on self-designed primer (Supplementary Table-1): Average Cq ± sme of w15-b320 (28.19 ± 7.91) against w15-btem (28.19 ± 6.21)). With TaqMan assay, the results were more homogeneous showing that supplementation with miR-320-p/DOSP indirectly silenced the miR-375-5p form (Figure 2E, F).

In the brain stem, the early-weaned females showed a down-regulation of *sert* and *drd1*, unlike males. Both male and female early-weaned rats displayed a deregulation of *cck* and *sert* (Figure 7, Supplementary Figure 5).

In Figure 7C representing hippocampus data, the miR-504 was down regulated for all groups when supplemented with miR-320-3p/DOSP indicating a strong effect of this miRNA supplementation. It should be underlined that no difference was found between sex and weaning times.

In summary, the supplementations with miR-320-3p/DOSP or miR-375-3p/DOSP were more potent with the young rats raised with regular weaning. Surprisingly, the early-weaned male rats were more resilient to miRNA treatment as their relative levels of miR-320-3p were already very high. The miR-504 was unchanged in hypothalamus, down regulated in hippocampus, but up regulated along with miR-320-3p and miR-16-5p in females treated by miR-320-3p/DOSP.

## Discussion

Oral supplementation by miRNA-320-3p or miR-375-3p during lactation has long-term miRNA-specific consequences on the endogenous levels of corresponding miRNAs with a strong tissue-dependent memory. The long-term effect of miR-375-3p was more limited, according to the fourfold lower number of predicted targets than with miR-320-3p. Combining miR-320-3p/DOSP with early weaning enhanced miR-320-3p and chromogranin A expression in the duodenum. In the hippocampus, the miR-504 was down-regulated for both sexes, but in the brain stem, up regulated only for females, along with miR-320-3p and miR-16-5p levels. In the hypothalamus, clock levels were up regulated for both sexes. In the miR-375-3p/DOSP group, the density of enteroendocrine duodenal cells increased. The long-term effect of miR-375-3p/DOSP was more limited, according to the fourfold lower number of predicted targets than with miR-320-3p (Table 1).

**Table 1.** List of significantly altered micro and messenger RNAs in the duodenum, brain stem, hippocampus, and hypothalamus of early-weaned transgenic rat pups treated by a single oral supplement of miR-320-3p and sacrificed at Day-45. Note that the miR-375-5p level of w15-b320 was down-regulated in comparison with w30-b320 (p=0.02), and w30-btem (p=0.04). No difference was found between genes assayed on the stomach.

| Tissue | miRNA |  |  | mRNA |  |  |  |  |  |  |
| --- | --- | --- | --- | --- | --- | --- | --- | --- | --- | --- |
|  | miR-320-3p | miR-504 | miR-16-5p | drd2 | sert | 5ht1b | cck | gfp | clock | per1 |
| Duodenum | Up |  |  |  |  |  |  |  |  |  |
| Brain stem-Female | Up | Up | Up |  | Down |  | Down |  |  |  |
| Brain stem-Male |  |  |  |  |  |  |  |  |  |  |
| Hippocampus-Female |  | Down |  |  |  |  |  |  |  |  |
| Hippocampus-Male |  | Down |  |  |  |  |  |  |  |  |
| Hypothalamus-Female |  |  |  |  |  | Up |  |  | Up | Up |
| Hypothalamus-Male |  |  |  | Up |  |  | Up | Up | Up |  |

The miRNA/DOSP complexes are delivered in the stomach, but according to the described kinetic^20^, they can also be delivered in proximal sites of the small intestine. Our transgenic rat model is allowing us to explore the influence of miRNA supplementation at distance from the inoculation site on the neuroendocrine cell lineage of the duodenum. Here we are using the miR-320-3p with cytoplasmic and nuclear sites of bioactivity, in parallel with miR-375-3p with bioactivity limited to the cytoplasm. Under our hand, the administration of DOSP loaded with a specific miRNA can be considered neutral for the physiological effects triggered by the miRNA. Even if, paromomycine, the polar headgroup of DOSP, has been re-evaluated as potentially targeting the mammalian ribosome machinery.^49^ Our current vector is by-passing the physiological sidt1-adsorption of miRNA in the stomach.^8^ Remarkably, combining force-feeding miR-320-3p/DOSP and early weaning stress did not alter sidt1 transcripts. Likewise, we did not detect the loading of miR-320-3p or miR-375-3p in gastric extracellular vesicles. The sequences of these miRNAs have no sorting sequences in exosomes^50^, this is in favor of the absence of re-export of these miRNAs after their cytoplasmic delivery either toward the gastric lumen or into the blood. No immediate effect of miR-320-3p/DOSP is described in plexus choroid nor cortex.^20^ In fact, our data are indicating a very high variability of miR-320-3p detection after 8 hours, in the plasma of miR-320-3p/DOSP or miR-375-3p/DOSP groups compared to controls. It seems highly likely that DOSP complexes were able of getting through the digestive epithelium into the plasma, then reaching the brain-blood barriers. We cannot label our DOSP vector for fluorescent tracking in blood and lymph but, DOSP could be tailored for targeting specific gut cell lineages, and for taking into account the putative interaction with the ribosome machinery. Such a vector would help resolving the paradoxe of a gut delivery with consequences in the brain area on the levels of endogenous miR-320-3p or miR-375-3p.

Control rats submitted to early weaning, had deep alterations in the levels of endogenous miR-320-3p in duodenum and of miR-320-3p and miR-375-3p, and all brain compartments tested (Figures 2, 3, 4, 5). These data are highly supportive of using miR-320-3p, present in breast milk, as supplements in breast-fed rat. Our data are indicating a down regulation of miR-375-3p as well as of miR-320-3p in the hippocampus of early weaned young rats in contradiction with the increased expression of miR-375-3p in the hippocampus of stressed mice.^32^

Our data are showing that in the brain but not in the gut, sex is playing a critical role, confirming McKibben et al (2021)^58^ who have found that in the hypothalamus, miR-132-3p and miR-504 are responsive to Early-Life stress, with males expressing greater changes following postnatal stress.

Surprisingly, our supplementation by miR-320-3p/DOSP has more impact on targeted miRNAs in the young rats raised with regular weaning as compared to early-weaned rats. The profile induced by miR-320-3p/DOSP supplementation is also driving the endogenous miR-375-3p regulation. By contrast, the effect of miR-375-3p/DOSP supplementation is weaker than with miR320-3p according to the limited subset of transcripts under the regulation of this miRNA. We are showing an increased density of enteroendocrine GFP-labelled cells, suggesting that the concentration of miR-375-3p were high enough to be delivered in duodenal proliferative or stem cells with late consequences on the kinetics of the duodenum. Likewise, the chromogranin A and GIP transcripts were, relatively to control bolus, decreased in young rats subsequently to miR-375-3p/DOSP and increased after miR-320-3p/DOSP. The level of endogenous miR-320-3p at Day-45 was up regulated in the duodenum (Figure 2B) indicating that the miR-320-3p/DOSP treatment can restore this miRNA level according to its relative expression level after regular weaning. Future works could explore the effects of miRNA supplementation directly in the stem cells of duodenal epithelia. New therapeutic for preventing early life stress may take advantage of the possibility to precisely target duodenal enteroendocrine cells.

A single bolus of miRNAs before weaning has induced in young rats, long-term effect on the expression of several miRNAs and mRNA, depending on the miRNA given by force-feeding. In the stomach, the levels of endogenous miR-320-3p and miR-375-3p (Figure 2 A) were significantly lower with rat pups treated by miR-320-3p/DOSP compared to controls and miR-375-3p/DOSP-treated rat pups. In all brain compartments tested, we have found that endogenous mir-320-3p and 375-3p were significantly up-regulated for rat pups treated with miR-320-3p/DOSP compared with controls and miR-375-3p/DOSP-treated rat pups both for female (Figure 4) and male (Figure 5).

Force-feeding rat pups with miR-375-3p is targeting fewer genes than miR-320-3p, consequently delivering miR-375-3p is without any effect on the endogenous level of miR-320-3p. Our data are in favor of a non-described hierarchical molecular link between endogenous miR-320-3p and miR-375-3p. Force-feeding rat pups with miR-320-3p/DOSP is revealing that, like predicted with our *in silico* data showing a wider target range for this miRNA, the rat pups with regular weaning have deregulation in the stomach as well as in brain compartments, some impacting endogenous miR-375-3p. In addition, we have detected the expression of miR-375-5p in duodenum extracts, without any difference between early-weaned rats treated by miR-320-3p/DOSP and controls. The miR-375-3p and 5p are both described in several rat tissue.^57^ Future research on a specific epithelial cell lineage is needed to explore the dynamic of the ratio between miR-375, 3 p and 5p molecules in single-cell. Young et al., 2022^60^ have shown that a stoechiometry exists between miR-140-5p and 140-3p with a physiological meaning for cartilage biosynthesis. Future works are needed in breast milk supplementation to take into account this risk of displacing the ratio between 5p and 3p for canonical miRNA like miR-375. To our knowledge, the incidence of a lower amount of miR-26a in mouse breast milk has been reported with physiological consequences in the adipocyte compartment (Pomar et al., 2021).^59^

The miR-320-3p is currently explored for its bioactivity in various diseases from type-2 diabetes to atherosclerosis.^26–28^ The *in vivo* delivery of miR-320-3p is targeting binding sites located both on polr3d promoter and on polr3d 3’UTR. Polr3d is the subunit-17 of polymerase-III involved in tumorigenesis. The RNA polymerase III is now considered linked to aging and longevity through its action on TORC and insulin genes as well as its activity on genes related to telomerase activity.^51, 52^ However, we have shown only an immediate effect on the chromatin complexes related to H3K4me3, as well as an absence of long-term effect on polr3d mRNA. The miR-320-3p has been studied for post-transcriptional gene silencing in the cytoplasm of rat endothelial and cardiac cell cultures derived from diabetes situations, on several genes among which the *heat shock protein family B (small) member 6*.^53^ The *hspb6* (also *hsp20*) gene is highly expressed in several organs including the stomach.^54^ Our data are confirming the immediate effects on *polr3d* and *hspb6* genes^20^, but additional works are needed to explore the long-term putative effects on polr3d complex which includes 17 subunits; as well as any effect on telomerase activity.

Early postnatal life is a critical period where stressful experiences may have the potential for long-term programming. The application of such preventive and therapeutic approaches during early-life sensitive periods is likely to be particularly promising. If one could modify the epigenetic patterns disrupted by exposure to stress through specific epigenome-targeted therapeutic interventions, then it would be possible to correct the impaired patterns of gene expression to prevent the stress-induced chronic pathologies and to improve human health and longevity. The early-weaned rats were more resilient to miR-320-3p/DOSP treatment on the expression of endogenous miR-320-3p and miR-375-3p. The innate immunity of early-weaned rats at the stomach level is also deeply altered, in part linked to the alteration in gastrointestinal permeability.^55^ However, our treatment with miR-320-3p/DOSP did not induce significant evolution of the cytokines related to immunity. Immune dysregulation is considered to be a key pathway linking the childhood adversity to elevated rates of morbidity and mortality from a number of chronic diseases later in life. Note that we did not report up-regulation of miR-375-3p related to the double stress of miR-320-3p/DOSP and early weaning. As shown in Figures 3 and 7, the networks of genes significantly deregulated in the stomach or brain compartments for early weaned rats are very narrow compared to the networks obtained after a regular weaning.

We have described the modification of clock transcripts in the hypothalamus or the liver of young Wistar rats at Day-35 showing an increased level in the nocturnal situation.^36^ Our data obtained on transgenic Sprague-Dawley rats have been obtained with a sacrifice done in the nocturnal phase (Figure 1). However, despite *in silico* prediction, the miR-320-3p, 16-5p and 132-3p were not correlated with *clock* transcripts (Figure 6 B-D). Further experiments are needed to explore whether the effect of early-weaning stress combined or not with force-feeding miR-320-3p alters the circadian clock machinery. In Figure 7B, with early-weaned rats force-fed with miR-320-3p, clock levels were high in the hypothalamus of males and period1 in females (Table 1). Future works in the developmental biology of the circadian clock could open an efficient therapeutic avenues.

In conclusion, our supplementation of lactating rat pups with extracellular miR-320-3p given before early weaning stress alters the miR-320-3p expression in duodenum, the miR-504 expression in the brain stem of female, and of clock transcripts in hypothalamus (Table 1), calling for behavioral studies. We are describing a new relationship between 2 unrelated miRNAs, miR-320-3p and miR-375-3p underlining a hierarchy between miRNA networks. The exploration of therapeutic potentials of miRNAs needs an approach in integrative physiology with a highly specific site of delivery like duodenal enteroendocrine cell lineage and articulated around the competing endogenous RNA hypothesis.^61^ This approach would gain much momentum by the implementation of results in an international database, improving the gap between *in silico* prediction and biological observations. The development of a new milk formulation intended to manipulate the epigenetics of the baby will benefit from such preclinical models.^62^

## Acknowledgments

We thank Mrs. Isabelle Grit and Mr. Alexis Gandon, both of UMR-1280, for help in rat care and sacrifice. We are also indebted to Dr. Patricia Parnet and Dr. Hervé Blottière (UMR-1280) for constant support. Part of these data have been presented during the PhD defense of Gabriel Tavares (Gabriel A Tavares, (2018-2021) Thèse en co-tutelle internationale, Université de Nantes (Bertrand Kaeffer), Université de Pernambuco (Sandra L de Souza) http://www.theses.fr/s276984) and for the Master-2 of Mrs Maïwenn Queignec. The authors would like to appreciate that some figures in this article were created with BioRender.com.

## Contribution of authors and correspondence

GT was in charge with brain dissection, handling, and transcript analyses. AT did *in silico* analysis, part of duodenum and stomach transcript analyses. BC, LB, SR, IA, GLD have constructed and maintained the transgenic rat strain. MQ and GLD realized immunohistology experiments. BP provided DOSP and know-how. BK, MQ, GT inoculated rat pups.

BK realized ChIp experiments, acting as coordinator of sacrifices, storage of samples, and centralizing analyses. All authors have contributed to the manuscript.

---

**Supplementary Table 1.**
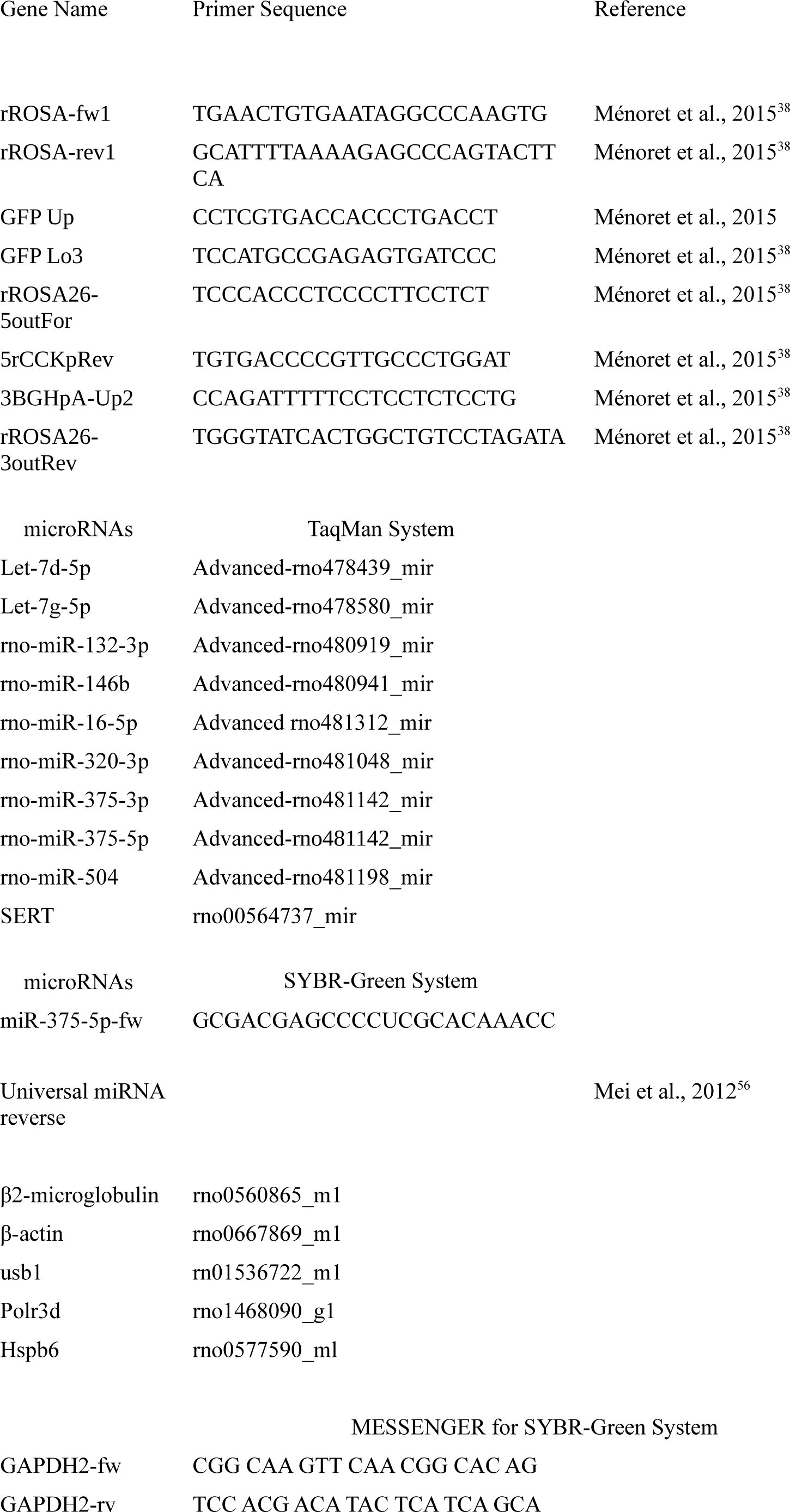
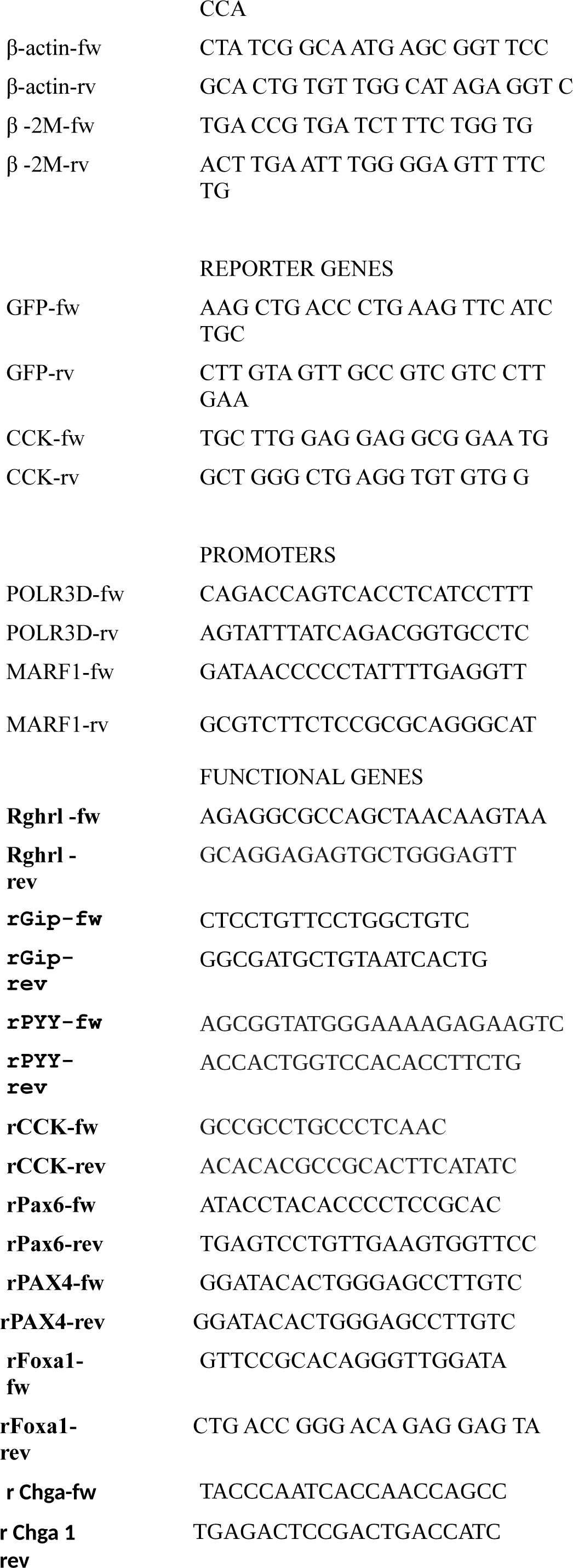
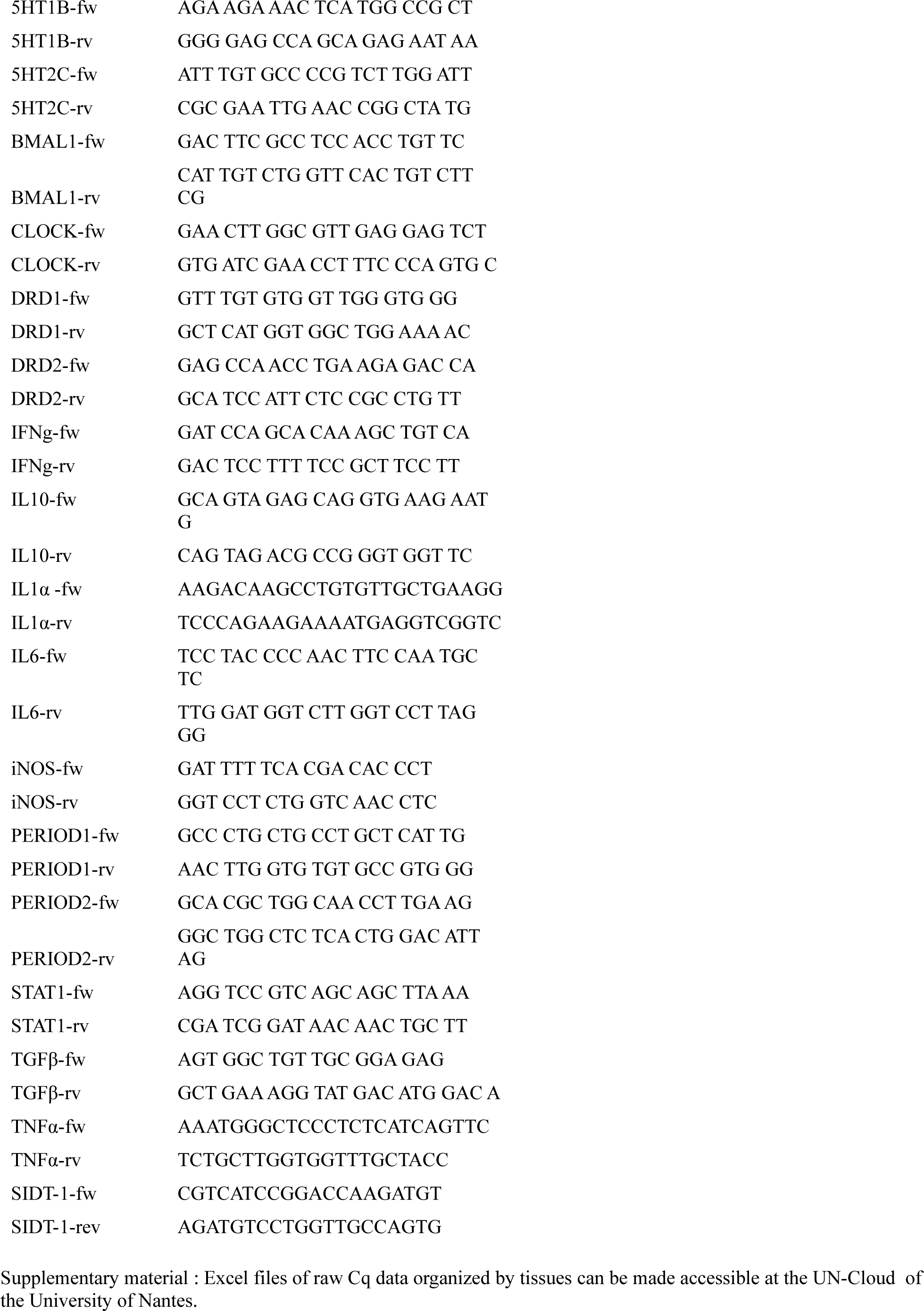
Details of the primers used to construct the transgenic rats, and quantified 6 miRNA and 42 mRNA.

**Supplementary Figure 1.**
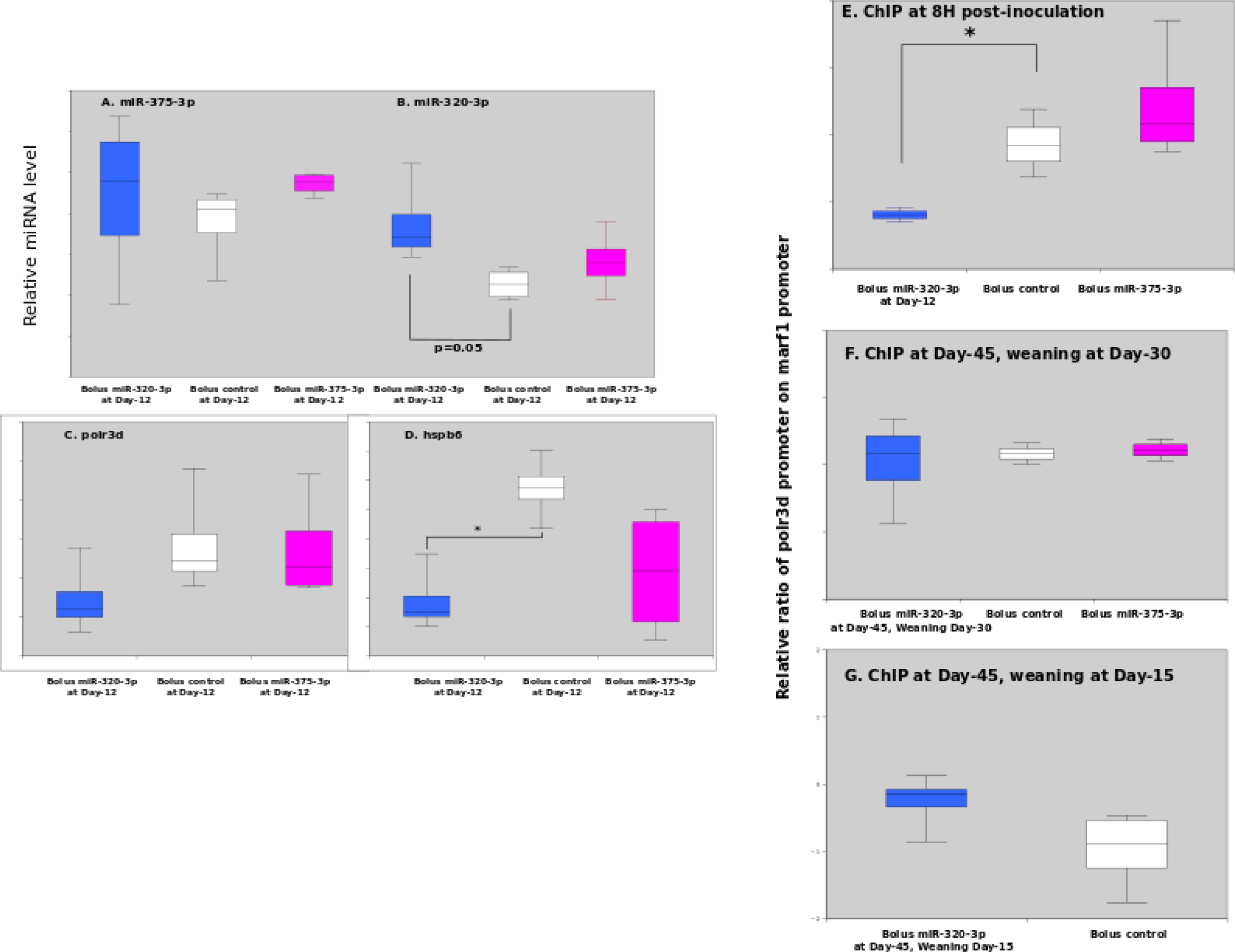
Immediate effect of miRNA supplementation. Evolution of miR-375-3p (A), 320-3p (B), polr3d (C) and hspb6 (D) mRNA, 8 hours after bolus for rat pups at Day-12 in stomach wall. Chromatin-immunoprecipitation assay against H3K4me3. Note the significant alteration at 8H after a bolus with miR-320-3p in gastric cells (E) and the absence of memory effect after regular (F) or early (G) weaning. The light gray background reminds that rats were sacrificed in the dark phase. Note: * p<0.05; ** p<0.01; *** p<0.001.

**Supplementary Figure 2.**
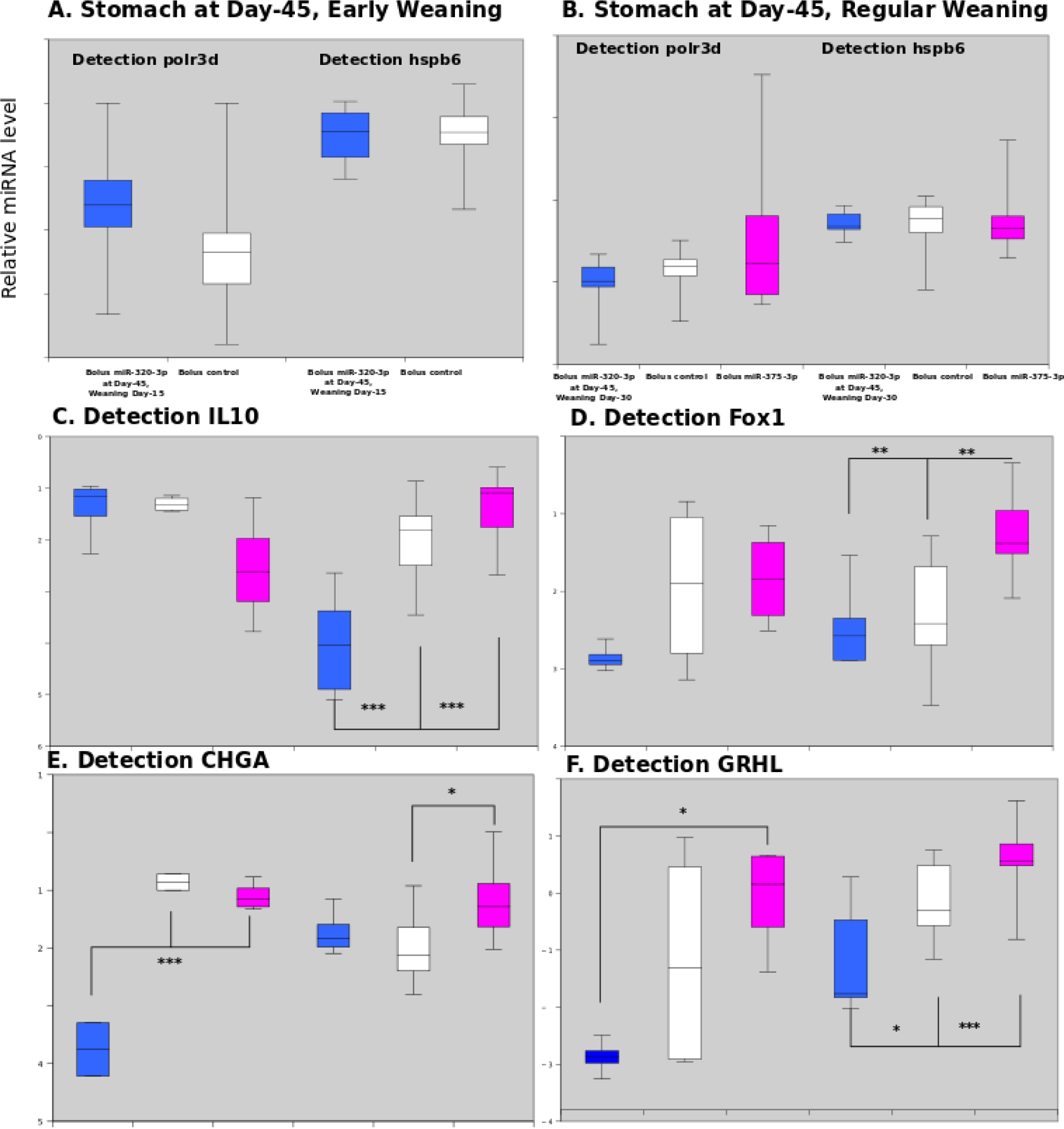
Evolution of polr3d and hspb6 mRNAs in stomach wall,at Day-45 after early (A) or regular (B) weaning. Concerning the inflammation status, the IL-10 [C], Fox1 (D), ChGRA (E), and GRHL (F) transcripts were all down-regulated at Day-45 for rat treated with miR-320-3p/DOSP. The light gray background reminds that rats were sacrificed in the dark phase. Note: * p<0.05; ** p<0.01; *** p<0.001.

**Supplementary Figure 3.**
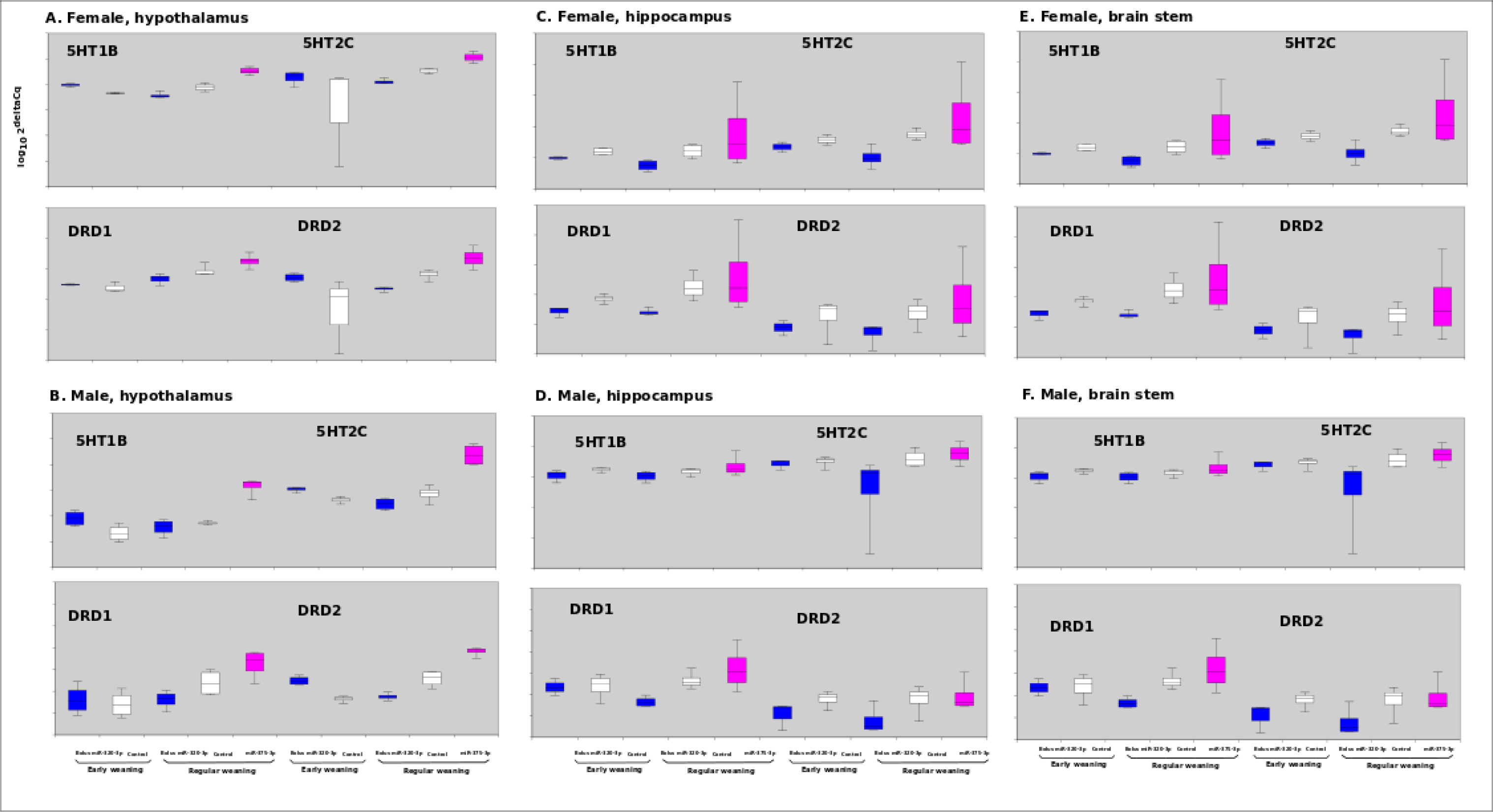
Evolution of transcripts related to serotonin and dopaminergic balance (5HT1B, 5HT2C, DRD1, DRD2) in hypothalamus (A, female; B, male), in hippocampus (C, female; D, male), and in brain stem (E, female, F, male) of rat treated with miR-320-3p or 375-3p/DOSP according to early or regular weaning. The light gray background reminds that rats were sacrificed in the dark phase. Note: * p<0.05; ** p<0.01; *** p<0.001.

**Supplementary Figure 4.**
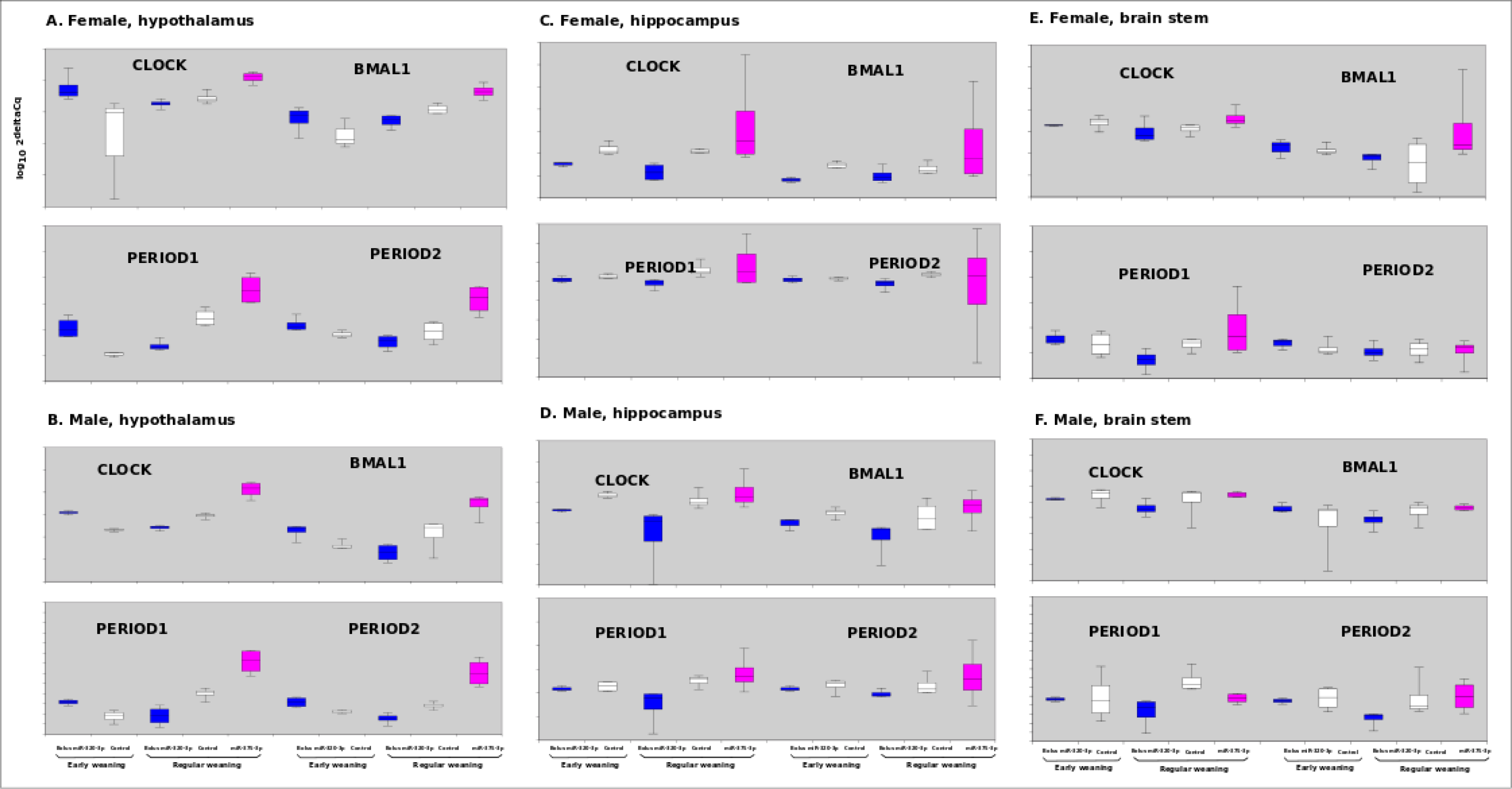
Evolution of transcripts related to the circadian clock (clock, bmal1, period1, period2) in hypothalamus (A, female; B, male), in hippocampus (C, female; D, male), and in brain stem (E, female, F, male) of rat treated with miR-320-3p or 375-3p/DOSP according to early or regular weaning. The light gray background reminds that rats were sacrificed in the dark phase. Note: * p<0.05; ** p<0.01; *** p<0.001.

**Supplementary Figure 5.**
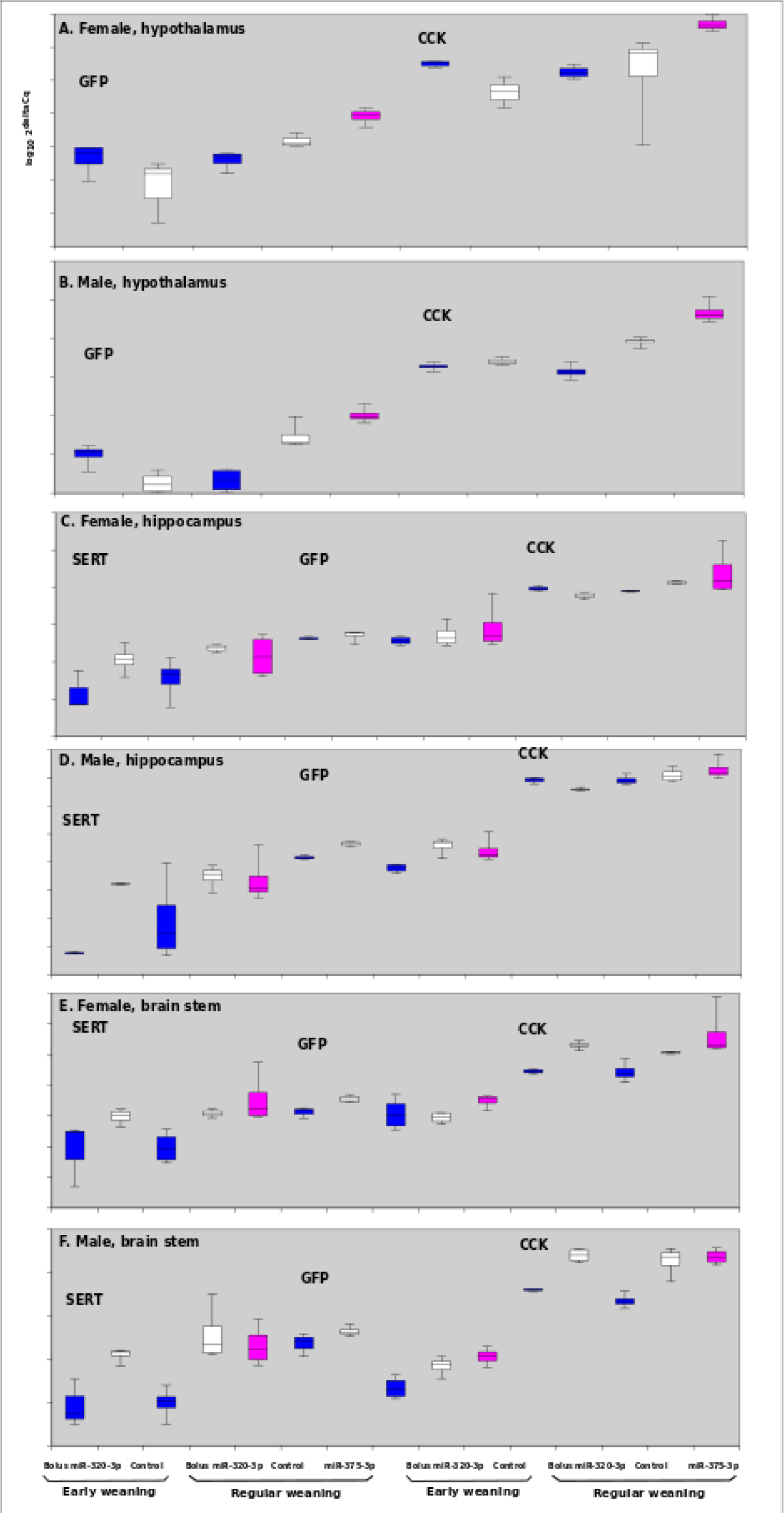
Evolution of SERT, GFP, CCK transcripts in hypothalamus (A, female; B, male), in hippocampus (C, female; D, male), and in brain stem (E, female, F, male) of rat treated with miR-320-3p or 375-3p/DOSP according to early or regular weaning. The light gray background reminds that rats were sacrificed in the dark phase. Note: * p<0.05; ** p<0.01; *** p<0.001.

